# Androgen Regulates SARS-CoV-2 Receptor Levels and Is Associated with Severe COVID-19 Symptoms in Men

**DOI:** 10.1101/2020.05.12.091082

**Authors:** Zaniar Ghazizadeh, Homa Majd, Mikayla Richter, Ryan Samuel, Seyedeh Maryam Zekavat, Hosseinali Asgharian, Sina Farahvashi, Ali Kalantari, Jonathan Ramirez, Hongyu Zhao, Pradeep Natarajan, Hani Goodarzi, Faranak Fattahi

## Abstract

Severe acute respiratory syndrome coronavirus 2 (SARS-CoV-2) infection has led to a global health crisis, and yet our understanding of the disease pathophysiology and potential treatment options remains limited. SARS-CoV-2 infection occurs through binding and internalization of the viral spike protein to angiotensin converting enzyme 2 (ACE2) on the host cell membrane. Lethal complications are caused by damage and failure of vital organs that express high levels of ACE2, including the lungs, the heart and the kidneys. Here, we established a high-throughput drug screening strategy to identify therapeutic candidates that reduce ACE2 levels in human embryonic stem cell (hESC) derived cardiac cells. Drug target analysis of validated hit compounds, including 5 alpha reductase inhibitors, revealed androgen signaling as a key modulator of ACE2 levels. Treatment with the 5 alpha reductase inhibitor dutasteride reduced ACE2 levels and internalization of recombinant spike receptor binding domain (Spike-RBD) in hESC-derived cardiac cells and human alveolar epithelial cells. Finally, clinical data on coronavirus disease 2019 (COVID-19) patients demonstrated that abnormal androgen states are significantly associated with severe disease complications and cardiac injury as measured by blood troponin T levels. These findings provide important insights on the mechanism of increased disease susceptibility in male COVID-19 patients and identify androgen receptor inhibition as a potential therapeutic strategy.

## Introduction

Coronavirus disease 2019 (COVID-19) caused by severe acute respiratory syndrome coronavirus 2 (SARS-CoV-2) has become a pandemic affecting millions of people worldwide. Limited understanding of the disease pathophysiology has impeded our ability to develop effective preventative and therapeutic strategies. Multi-organ failure is the most lethal complication of SARS-CoV-2 infection^1^. Early reports from China show higher mortality rates in patients that develop myocardial injury, as measured by the elevation of serum troponin T levels^2,3^. Furthermore, COVID-19 patients presenting with cardiac injury were more likely to have a baseline cardiovascular disease such as hypertensive disorders or heart failure^2^.

It has been well documented that organ involvement and disease manifestation are correlated with the expression of SARS-CoV receptor and co-receptors on the membrane of target cells^4^. The spike (S) protein, which is responsible for the characteristic crown-like shape of coronaviruses, facilitates the binding of the virus to its receptors on the cell membrane^5,6^. ACE2 has been identified as the main receptor utilized by SARS-CoV-2 and SARS-CoV-1 to enter cells^7^. Additionally, recent findings using human and animal cell lines have demonstrated that SARS-CoV-2 relies on the serine protease TMPRSS2 for S protein priming prior to ACE2 facilitated internalization^6^. Research after the SARS-CoV-1 epidemic shows that knocking-out ACE2 results in markedly decreased viral entry in lungs of mice infected with the virus^8^. Recently, Tucker et. al. showed expression of ACE2 in human adult cardiac cells, such as cardiomyocytes, fibroblasts, pericytes, and vascular smooth muscle. Interestingly, ACE2 levels in cardiomyocytes are found to be increased in patients with failing hearts, further linking the expression profile of ACE2 to clinical findings in COVID-19 patients^9^. In this study, we identified pharmacological strategies to reduce the levels of ACE2 in human embryonic stem cell (hESC) derived cardiac cells in order to facilitate therapeutic interventions aimed at reducing viral entry, thereby mitigating the multi-organ complications induced by SARS-CoV-2 infection.

To identify candidate drugs capable of modulating ACE2 protein levels, we took advantage of our previously established methods to generate cardiac cells from human embryonic stem cells at large scale^10–12^ and performed a high content screen with a library of 1443 FDA-approved drugs and a subsequent *in silico* screen with the ZINC15 library of over 9 million drug-like compounds. We discovered that drugs most effective in reducing ACE2 protein levels converge on androgen receptor (AR) signaling inhibition as a common mechanism of action. These drugs were effective in reducing ACE2 and TMPRSS2 levels on the cellular membrane and resulted in reduced internalization of SARS-CoV-2 spike-RBD in hESC-derived cardiac cells and human primary alveolar epithelial cells.

Clinical case studies have identified male sex as a major risk factor for SARS-CoV-2 complications. In fact, 70% of the patients on ventilators in the ICU were found to be males^1^. To explore the possible role of androgen signaling on poor disease outcomes in male COVID-19 patients, we conducted a study on two independent cohorts of patients tested for SARS-CoV-2. Among males, we found a significant positive association between free androgen index and the risk of severe COVID-19 and between prostatic disease as a surrogate for androgen dysregulation and abnormal blood troponin T levels. Our data provide a potential mechanistic link between clinical observations and pathways involved in COVID-19 pathogenesis. The results identify AR signaling inhibition as a potential therapeutic strategy to reduce SARS-CoV-2 viral entry and mitigate severe manifestations in COVID-19 patients.

### High-throughput drug screen identifies ACE2 modulators in human cardiac cells

Our analysis of previously published single cell RNA sequencing datasets^13–15^ showed abundant expression of SARS-CoV-2 receptor, ACE2, and co-receptors, TMPRSS2 and FURIN in adult cardiac, esophageal, lung and colon tissues (Supplementary Figure 1A). Given the highly significant association of poor outcomes in COVID-19 patients with cardiovascular complications^2,3^ and the significant role of ACE2 in cardiac physiology^16^, we chose to focus on the regulation of ACE2 levels in cardiac cells. Due to limitations associated with isolation and maintenance of human cells from primary tissue, we used our previously established hESC differentiation method as an alternative strategy to generate cardiac cells (Supplementary Figure 1B)^12^ Previously published transcriptomics data on hESC-derived cells generated using this method^12^ confirmed the expression of SARS-CoV-2 receptor and co-receptors mRNAs in cardiomyocytes and non-cardiomyocyte (Supplementary Figure 1C). The differentiated cells also stained positive for ACE2 as assessed by immunofluorescence imaging (Supplementary Figure 1D).

Searching for modulators of ACE2 levels in hESC-derived cardiac cells, we screened a Selleckchem small molecule library composed of 1443 FDA-approved drugs (Figure 1A). ACE2 levels were measured in drug-treated cells using high content imaging and a list of drugs that significantly down-regulate or up-regulate ACE2 were identified based on their normalized ACE2 expression z-scores (Figure 1B, Supplementary Table 1). We confirmed the effect of a compound with a high positive z-score (vincristine) and a compound with a low negative z-score (dronedarone) on ACE2 fluorescence intensity (Figure 1C), and subsequently selected several hit compounds with low and high z-scores (Supplementary Figure 1 E&,F) for further analysis and validation at 1 μM and 2 μM concentrations (Figure 1D-E). The high-quality cell-based measurements and the inherent diversity of the FDA library provided a unique opportunity to develop a virtual high-throughput screening (vHTS) approach that allowed for rapid *in silico* screening and costeffective identification of compounds that can elicit the desired biological response. Combined analysis of these *in vitro* measurements and *in silico* predictions allows us to nominate molecular entities that can effectively modulate the signaling pathways responsible for ACE regulation.

**Figure 1.**
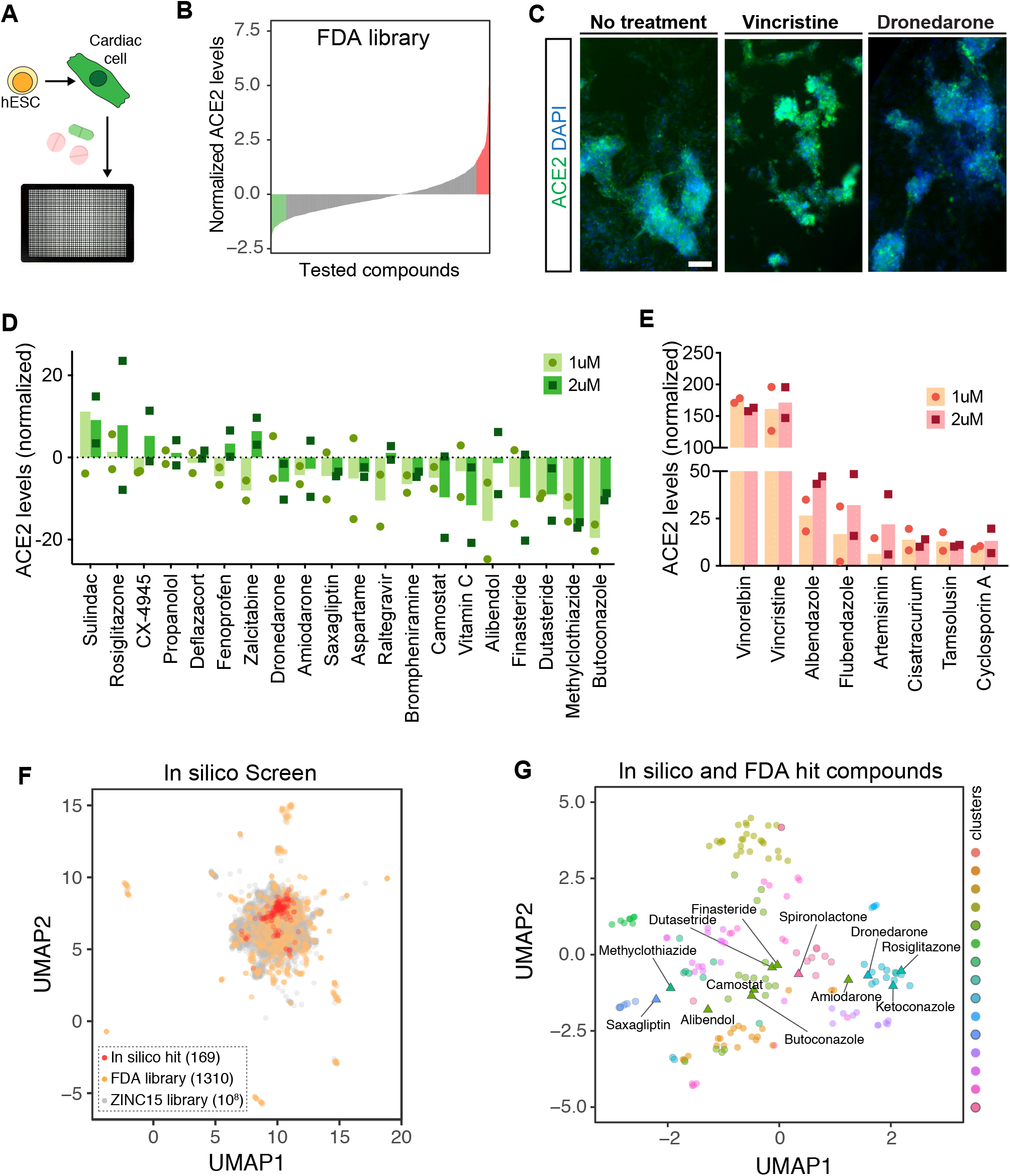
High-throughput *in-vitro* and *in silico* screenings identify drugs that modulate ACE2 expression in hESC-derived cardiomyocytes. **A-B)** High-throughput screening of Selleckchem FDA-approved drug library identifies drugs that increase and decrease ACE2 expression in hESC-derived cardiomyocytes. **C)** Representative immunofluorescent images of cells treated with vehicle, vincristine and dronedarone at 1μM. Scale bar:100 μm. Dose response of hits that **D)** decreased, and **E)** increased ACE2 expression in hESC-derived cardiomyocytes culture. F) two dimensional visualization of molecular features (Morgan fingerprints) for the *in vitro* and *in silico* tested compounds using UMAP. For the ZINC15 library, the points are sub-sampled by a factor of 10^3^. G) UMAP visualization of the *in vitro* (labeled) and *in silico* (unlabeled) hit compounds. Also shown are the K-means cluster memberships based on their Morgan fingerprints.

To achieve this goal, we first randomly split our *in vitro* screening results into three datasets as training, validation and test inputs (Supplementary Figure 2A-C). We used the model trained on the *in vitro* screening data to predict changes in ACE2 expression in response to treatments with the *in silico* library. We then variance normalized the model predictions to determine their associated *z*-scores. We selected the compounds with a z-score smaller than −4 as “hit” compounds. To visualize these hits, we used Morgan fingerprints to generate molecular features for all *in vitro* and *in silico* tested compounds and visualized their relationship in a 2-dimensional space using Uniform Manifold Approximation and Projection (UMAP) (Figure 1F). We then used K-means clustering to categorize *in silico* and *in vitro* hits and grouped them into 15 coarse-grained clusters based on their molecular features (Figure 1G). We observed a consistent coclustering of the FDA-approved drugs that are known to have structural similarities, such as the cluster containing finasteride and dutasteride, confirming the utility of these clustering representations. Taken together, this integrative cell-based and *in silico* screening strategy enabled the identification and nomination of novel drug-like compounds with similar structure and potentially higher potency as compared to the FDA-approved drugs utilized in the *in vitro* screen.

### Drugs that reduce ACE-2 regulate steroid signaling and peptidase activities

We explored if a potential shared pathway exists among the candidates that decrease ACE2 levels on the membrane of cardiac cells. We acquired isometric simplified molecular-input line-entry system (SMILES) for each drug in the FDA-approved library from Selleckchem and used them to predict drug-protein interactions via the similarity ensemble approach (SEA) computational tool^17^. The SEA-predicted drug-protein pairs were filtered, selecting human proteins and predicted interaction *p*-values <0.05, which yielded 2150 predicted proteins targeted by the drug library.

Weighted combined z-scores were then calculated by adding normalized *z*-scores across all compounds that target each target protein^18^. The *p*-values were then calculated based on the combined z-scores and adjusted using p.adjust (method=false discovery rate(FDR)). As an orthogonal approach, for each protein, we recorded the number of treatments with negative normalized *z*-scores as well as the total number of compounds predicted to target that protein. Using the sum of counts for all other targets and drugs, we performed a Fisher’s exact test to evaluate the degree to which negative *z*-scores were enriched among the drugs likely to target a protein of interest. As expected, the two *p*-values, i.e. combined *z*-score and Fisher’s, are generally correlated (R=0.6, *p*<1e-200). In Figure 2A, we have specifically compared these *p*-values across the genes with negative average *z*-scores. Finally, we selected the likely target genes using the following criteria: average *z*-score<0, FDR<0.25 based on combined *z*-score analysis, and Fisher’s *p*<0.05. This selection process resulted in 30 proteins nominated as significant drug targets (Figure 2B,C, Supplementary Table 2).

**Figure 2.**
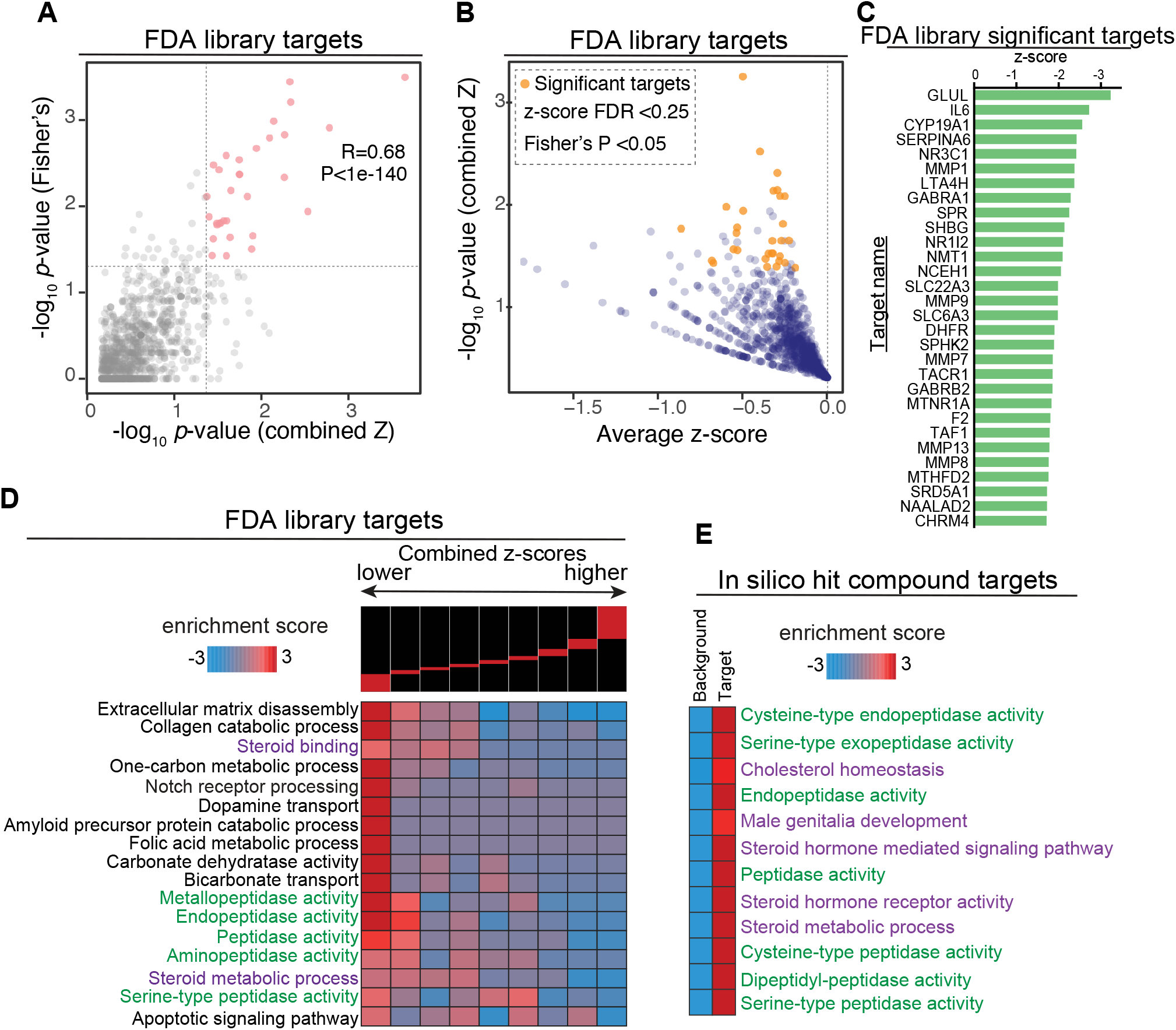
Target prediction analysis identified shared pathways among ACE2 regulators. **A)** We employed two independent tests for identifying the genes that are most likely targeted by the effective treatments: (i) a combined *z*-score approach, where normalized *z*-scores from all the treatments associated with a gene are integrated, and (ii) a Fisher’s exact test to assess the enrichment of a gene among those that are targets of treatments with negative *z*-scores. Here we have shown the correlation between the *p*--values reported by these two independent approaches. **B)** A one-sided volcano plot showing the average *z*-score vs. −log of *p*-value for all genes with negative *z*-scores. The genes that pass our statistical thresholds are marked in gold (combined *z*-score FDR<0.25 and Fisher’s *p*-value <0.05). **C)** The identified target genes along with their combined *z*-score, associated *p*-value and FDRs. Also included are the total number of compounds each gene is likely targeted by, and the number of those that result in lower ACE2 expression (*z*-score<0). **D)** Gene-set enrichment analysis using iPAGE for the target genes identified from FDA-approved library with negative *z*-score. Genes were ordered based on their combined *z*-score from left to right and divided into nine equally populated bins. The enrichment and depletion pattern of various genesets is then assessed across this spectrum using mutual information. Red boxes show enrichment and blue boxes show depletion. **E)** Gene-set enrichment analysis for the *in silico* hits. Similar to (D), genes were grouped into those that are likely targeted by the identified compounds and those that are not (i.e. background). We then assessed the enrichment of each pre-compiled gene-set among the targets using iPAGE.

The previously published bulk transcriptomics data^12^ confirmed that 28 of these 30 proteins were expressed by the *in vitro* generated cardiomyocytes and/or non-cardiomyocytes supporting their potential as drug targets (Supplementary Figure 3A). Analysis of the collective expression of all 30 predicted targets using module scoring revealed that some ACE2 expressing cell types, such as cardiac fibroblasts and ciliated cells in the lung, express high levels of multiple predicted drug targets (Supplementary Figure 3B). Additionally, expression of the predicted targets was detected in ACE2 expressing cell types of the adult human heart, and the majority were also detected in ACE2 expressing cell types of other commonly affected organs (Supplementary Figure 4). This analysis also provides insight into non-ACE2 expressing cell types that may be affected by treatment with the hit compounds. In particular, tissue resident immune populations of the myeloid lineage in the heart, esophagus and lung also collectively express many of the predicted target genes (Supplementary Figure 3B,4). This is particularly interesting given the recent reports that myeloid lineage and epithelial cells are affected in severe SARS-CoV-2 infection cases^19^.

To identify biological pathways associated with changes in ACE2 expression, we used the combined *z*-scores across all proteins using iPAGE gene ontology (GO) analysis (Supplementary Figure 5). Interestingly, target proteins associated with compounds that reduce ACE2 expression were associated with various GO terms related to peptidase activity and steroid metabolic processes (Figure 2D). These pathways were then validated with the SEA predicted protein targets from the *in silico* library screen. Interestingly, hit compounds identified in the *in silico* screen also showed enrichment for targets involved in steroid hormone activity, steroid metabolic processes, and peptidase activity (Figure 2E). Given the strong clinical evidence on COVID-19 disproportionately affecting men, the well-established role of peptidase in modulating ACE2 and SARS-Co-V-2 co-receptors, and the possible link between ACE2 expression and sex hormones, we sought to uncover the specific drug-protein interactions driving enrichment of steroid and peptidase pathways. We first mapped the drug-protein interactions for the full list of predicted targets and compounds with z-score<-1.5 (Figure 3A). This revealed a list of 48 compounds with significant interactions with target proteins (Supplementary Figure 6). Next we created drug-protein matrices on proteins in “steroid metabolic process” and “serine-type peptidase” activity GO terms to map the interaction of drugs with corresponding targets (Supplementary Figure 7A,B). This analysis highlighted the interaction of several drugs including ketoconazole, spironolactone, finasteride and dutasteride with androgen signaling modulators such as SRD5A1 and SHBG (Figure 3A, Supplementary Figure 7A,B) whereas other drugs from the *in vitro* screen, such as camostat mesilate, sotagliflozin, and guanabenz acetate, are specific to the serine-type peptidase activity pathway (Supplementary Figure 7B).

**Figure 3.**
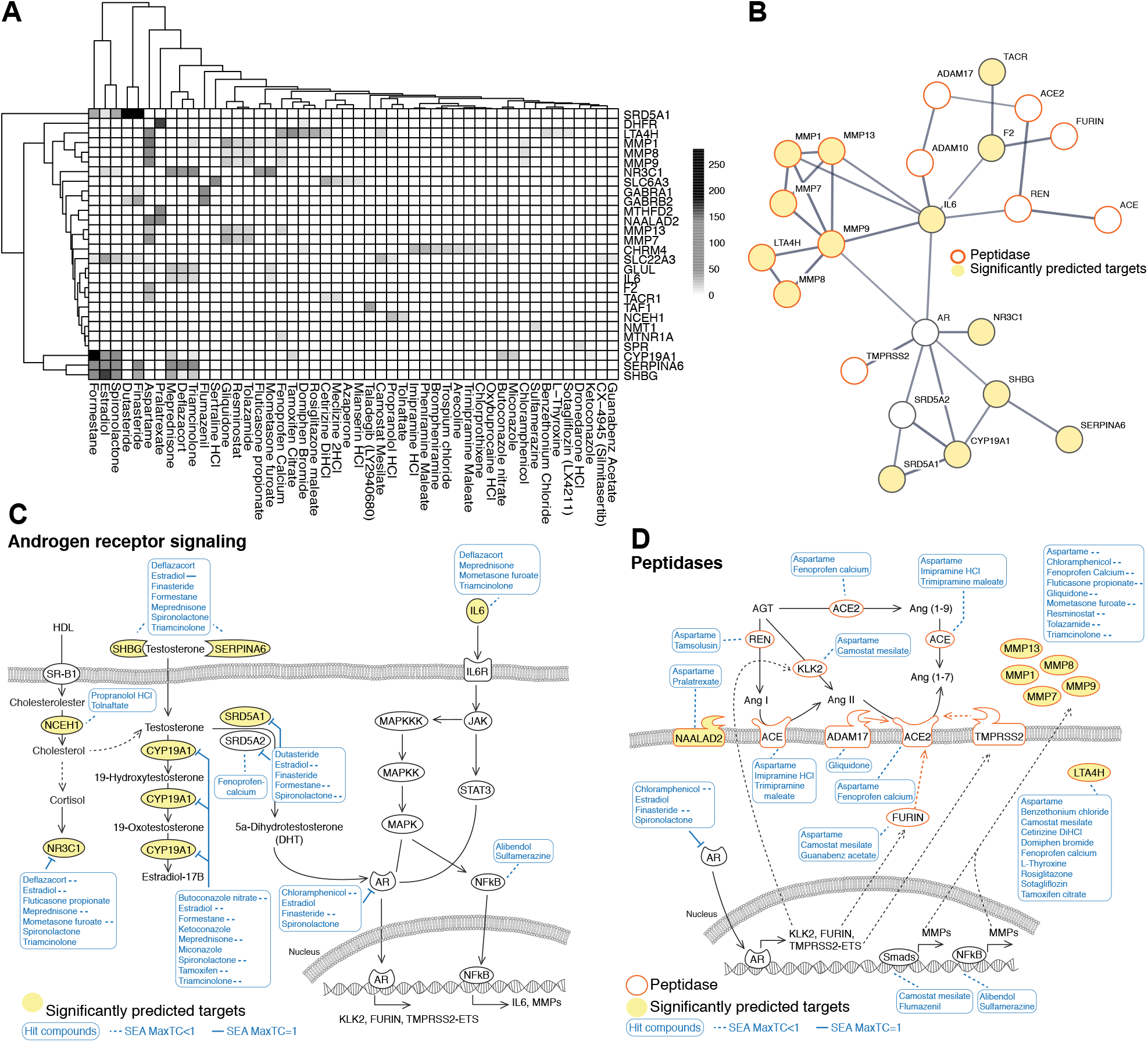
Androgen signaling regulates peptidase expression. **A)** The drug-gene interaction matrix for the 30 significantly enriched drug target genes from Figu 2C that are deemed functional in their respective analyses. Shading represents the significance of the predicted interaction. **B)** STRING protein-protein interaction network was used to identify interactions between our list of significantly enriched genes from Figure 2C (depicted as significantly predicted targets and yellow circles), androgen signaling pathway components (AR and SRD5A2), and proteins implicated in ACE2 regulation (ACE, ADAM10, ADAM17, FURIN, REN, TMPRSS2). Minimum required interaction score was set to 0.7 corresponding to high confidence and edge thickness indicating the degree of data support. **C)** SEA predicted drug-protein target interactions (blue lines and boxes) in the androgen signaling pathway. Yellow ovals represent significantly enriched genes from Figure 2C. Dashed lines represent MacTC <1. **D)** The expression of ACE2 related peptidases is regulated by AR and other transcription factors that are targets of our candidate drugs. MaxTC, maximum tanimoto similarity between compounds from ref_target to compounds from query_target in [0,1] with 1 being identical upto the resolution of the fingerprint.

To build a working model of the drug-protein interactions in ACE2 regulation, we conducted a proteinprotein interaction (PPI) network analysis to identify interactions between our list of significant predicted targets (Figure 2C), androgen signaling pathway components (AR and SRD5A2), and proteins implicated in ACE2 function regulation (ACE, ADAM10, ADAM17, FURIN, REN, TMPRSS2). We used the STRING physical interaction database to draw a high confidence network of associating proteins (Figure 3B). In the resulting network, AR and IL6 have the highest degree centrality, connecting to seven other nodes in the network. Furthermore, AR and IL6 share high betweenness centrality, connecting the androgen pathway module to the peptidases. The observed link between AR and IL6 is clinically important given the elevated IL6 response in severe cases of COVID-19 infection^20^.

The remarkable convergence of the gene set enrichment analysis and the PPI network on steroid hormone related genes and pathways prompted us to hypothesize that the drug candidates may be reducing ACE2 expression via inhibition of AR signaling and peptidase pathways (Figure 3C-D). Seven of the predicted drug targets are upstream regulators of AR signaling and are targeted by multiple drug candidates that reduce ACE2 (Figure 3C). Furthermore, peptidases such as FURIN and TMPRSS2 which are important players in ACE2 regulation and thereby SARS-CoV-2 viral entry^21,22^, are amongst the downstream targets of AR. These receptors and their upstream regulators are also predicted to be targeted by the drug candidates (Figure 3D).

### Androgen receptor signaling modulates ACE2 and TMPRSS2

Although most highly expressed in male reproductive organs, AR mediates hormone signaling in many male and female tissues^23^. In agreement, expression of AR and testosterone converting enzymes SRD5A1 and SRD5A2 are detected in ACE2 expressing cell types in the adult heart, lung, esophagus and colon (Supplementary Figure 8). Additionally, collective expression of genes involved in AR signaling (GO:0030521), common receptors upstream of AR activation^24–27^, and common gene targets of AR transcription factor activity^28^, reveals potential organ and cell type specific differences in AR signaling regulation (Supplementary Figure 8). A list of genes included in each module is provided in Supplementary Table 3.

To determine whether AR regulates the expression of SARS-CoV-2 receptors directly, we used an existing AR ChIP-seq dataset generated in LNCaP cells^29^ to identify direct transcriptional targets. The genes with AR binding to this 5kb downstream or upstream of transcription start sites (TSS) were selected as direct AR-bound targets. We next used a transcriptomics dataset generated using RNAi-mediated knockdown of AR^30^ and compared gene expression changes in response to AR knockdown. We divided log-fold expression changes into nine equally populated bins, which were also shown along with the patterns of AR enrichment and depletion at the corresponding TSS across the data (Figure 4A). This analysis identified ACE2, and other SARS-CoV-2 co-receptors TMPRSS2 and Furin as direct transcriptional targets that are downregulated in response to AR knockdown (Figure 4A). Furthermore, after exposing hESC-derived cardiac cells to the drug candidates that are known to inhibit AR signaling, we observed significant reductions in ACE2 and TMPRSS2 protein levels (Figure 4B,C). Dutasteride and spironolactone, the top candidates that modulate AR signaling, were both able to reduce ACE2 levels in a dose-dependent manner (Supplementary Figure 9 A,B). Together, these results indicate that AR signaling regulates expression of SARS-CoV-2 receptors.

**Figure 4.**
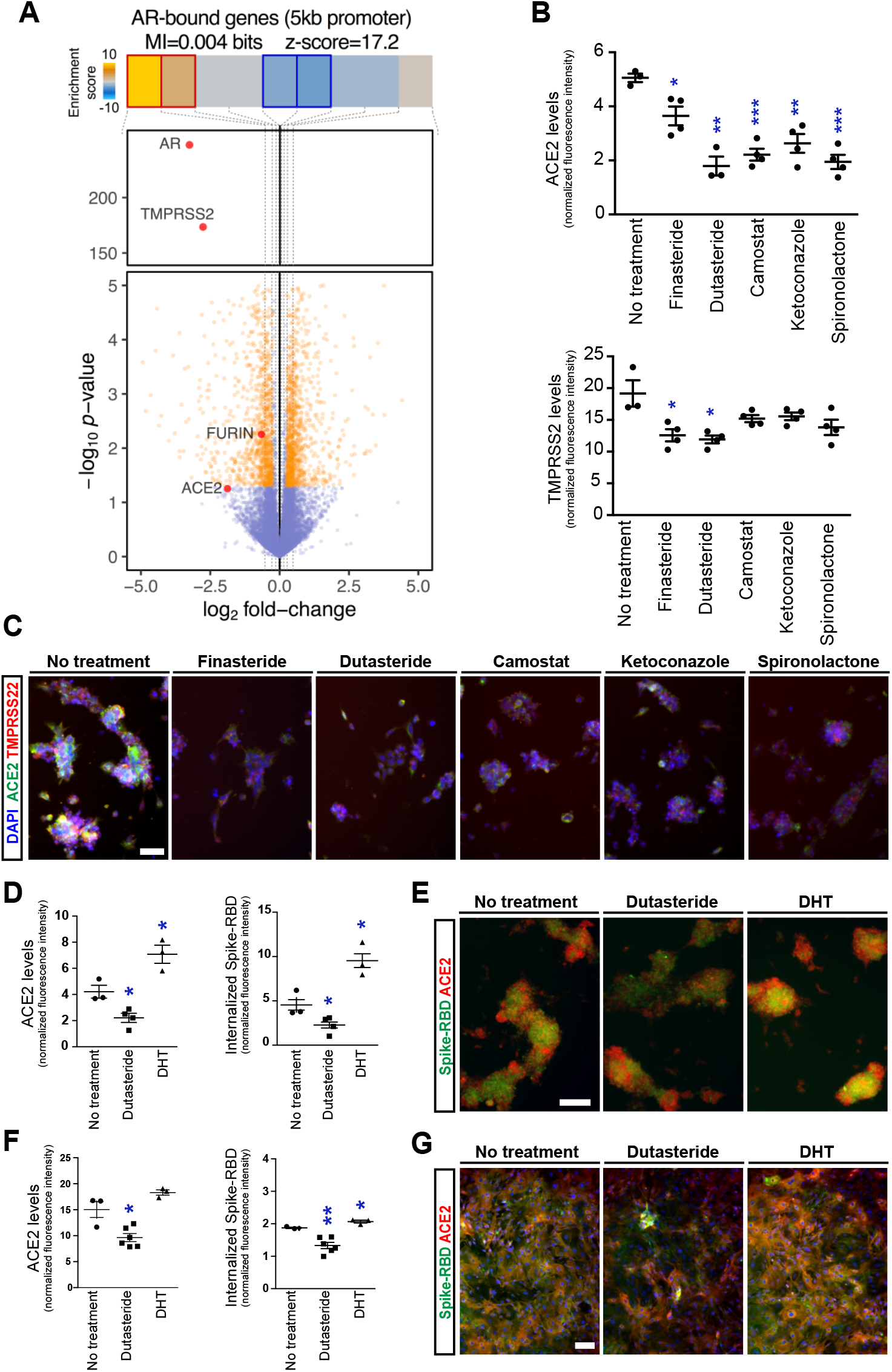
Androgen receptor signaling modulates ACE2 and TMPRSS2 levels and Spike-RBD internalization. **A)** Volcano plot visualizing gene expression changes in response to AR knockdown in LNCaP cells. Genes of interest are labeled and shown in red. Also shown are the enrichment and depletion pattern of AR target genes (i.e. genes with promoter AR binding) as a heatmap along with the mutual information value and its associated z-score. The log-fold change values were divided into equally populated bins and the enrichment of AR-bound genes in each bin was assessed using hypergeometric *p*-values and colored accordingly (gold for enrichment and blue for depletion; red and blue borders mark bins that are statistically significant. **B-C)** Effect of candidate androgen signaling inhibitors on ACE2 and TMPRSS2 levels on the surface of cardiac cells with their corresponding Immunofluorescence images. **D-G)** Differential effect of Dutasteride (potent inhibitor of Testosterone to DHT conversion) and DHT on the membrane ACE2 levels and spike-RBD protein entry to cardiomyocytes (D-E) and Pulmonary epithelial cells (F-G) with their corresponding immunofluorescence images. Scale bar= 50 μm in C and 100 μm in E &G.\**p*-value<0.05 \*\**p*-value<0.01 \*\*\**p*-value<0.001

To assess whether pharmacologic inhibition of AR signaling can reduce viral entry, hESC-derived cardiac cells were exposed to recombinant spike-RBD protein. After a 24 hour exposure, immunofluorescence imaging showed co-localization of spike-RBD protein with cells expressing ACE2 protein (Figure 4D-E). This supports the findings by Sharma et al. that hPSC-derived cardiomyocytes can be infected with SARS-CoV-2 virus^31^. Remarkably, pre-incubation with the AR signaling inhibitor, dutasteride, significantly reduced the levels of ACE2 and internalized spike-RBD, whereas, pre-incubation with the AR ligand and agonist, 5a-dihydrotestosterone (DHT), had the opposite effect. (Figure 4D-E). The effect of dutasteride on reducing ACE2 and spike-RBD internalization was further confirmed in human primary alveolar epithelial cells (Figure 4F-G).

### Androgen imbalance states are associated with COVID-19 complications in male patients

Our results suggest that androgen regulation can increase viral receptor and co-receptor expression leading to increased Spike-RBD entry. To determine whether androgen plays a role in COVID-19 disease manifestation, we explored the effect of disorders related to androgen imbalance on COVID-19 induced cardiac injury, measured by elevated troponin T levels. We also included the previously described risk factors associated with organ failure^32^ such as age, BMI, diabetes and hypertension in our data collection (Figure 5A). In the de-identified aggregate data from Yale New Haven Hospital, 1577 individuals tested positive for COVID-19 and had serum troponin T measured during the same encounter. There was a larger number of males with abnormal serum troponin T levels in both selected age groups (Supplementary Figure 10A). Association analysis in the COVID-19 patients showed that most risk factors are correlated with abnormal levels of troponin T in serum, but the risk factors also correlate with each other (Supplementary Figure 10B). To account for associations between individual risk factors, we tested multiple multi-variate models (as described in the methods section). In our final model, prostatic disease increased the odds of having abnormal troponin T by 50.5% (OR=1.505, 95% CI, *p*-value 0.046), independent of the other risk factors (Figure 5B).

**Figure 5.**
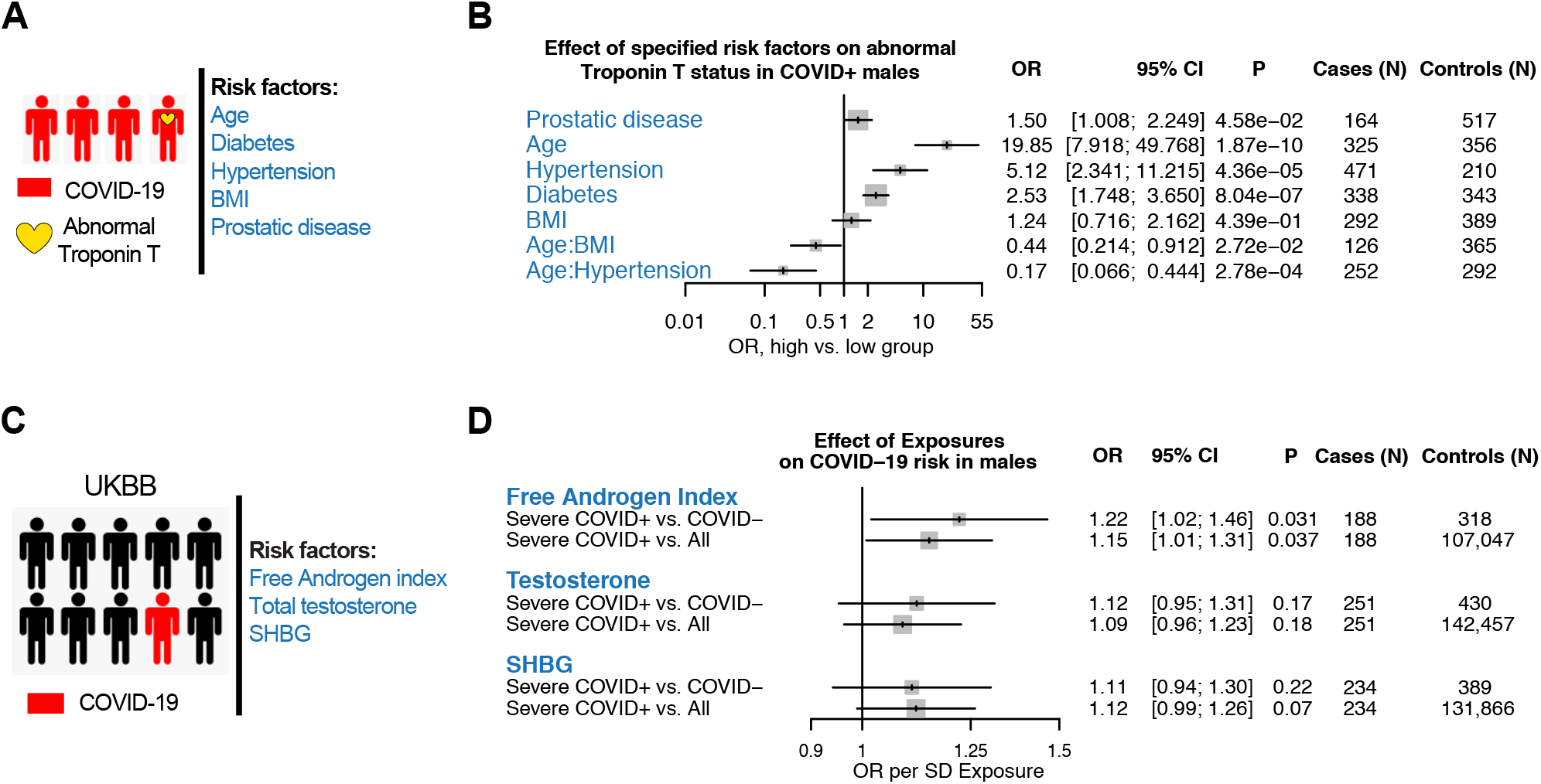
Effects of Androgen signaling on outcomes associated with COVID-19. **A.** Schematic representation of outcomes and risk factors studied in patients with Covid-19 at Yale New Haven Hospital. **B.** The effects of age, BMI, prostatic disease, hypertension and diabetes on the odds of having abnormal troponin T in male patients with Covid-19 in Yale patients. Age, troponin T and BMI were dichotomized during data collection. Age: 40-65y vs 65+, BMI: <30 vs >=30, troponin T: normal (<0.01 ng/ml) vs abnormal (>=0.01 ng/ml). For the primary outcome, the odds ratio were calculated for the pre-specified subgroups. **C.** Schematic representation of the outcomes and risk factors studied in the UK Biobank (UKBB) cohort. **D.** Association of androgen indices on Covid-19 susceptibility and severity in males among the UKBB. Association of normalized free androgen index, testosterone, and SHBG indices with two COVID-19 case/control models: for individuals tested with COVID comparing Severe COVID+ cases vs. COVIDcontrols, and for all white British individuals from the UK Biobank English recruitment centers with available COVID-test reporting, comparing Severe COVID+ cases vs. All controls. Severe COVID+ is defined as hospitalized with at least one COVID-19+ test. Analyses were adjusted for age, normalized Townsend deprivation index, normalized body mass index, and the first ten principal components of genetic ancestry. SHBG= sex-hormone binding globulin, Severe COVID+ = Tested positive for coronavirus disease 2019 and hospitalized, COVID-= Tested negative for coronavirus disease 2019, All = all individuals with reported COVID-19 testing from the UK Biobank English recruitment centers, OR = odds ratio, SD = standard deviation, CI= confidence interval.

To further test the effect of increased androgen on disease manifestations, we analyzed the association between serum androgen levels with disease severity in the UK Biobank (UKBB) (Figure 5C). A total of 311,857 individuals in the UKBB passed quality control criteria and were among those with reported COVID-19 testing from the UKBB English recruitment centers. 1,407 individuals were tested for COVID-19, of which 543 (37.5%) had at least one positive test. Of all the individuals who tested positive for COVID-19, 454 individuals (83.6%) were hospitalized, with evidence from their microbiological record of inpatient status (i.e. severe COVID-19; Supplementary Figure 11A). Quantitative measures for testosterone, sex-hormone binding globulin (SHBG), and free androgen index were first standardized to account for sex differences (Supplementary Figure 11B). Free androgen index was significantly associated with COVID-19 susceptibility and severity in males (Figure 5D), with no significant effect in females (Supplementary Figure S12), in both adjusted and unadjusted models. In particular, among males who were tested for COVID-19, each standard deviation (SD) increase in free androgen index increased the risk of a positive test (OR=1.22, 95% CI: 1.03-1.45, *p*-value 0.024), as well as severe COVID-19 infection (OR=1.22, 95% CI: 1.02-1.46, *p*-value 0.031), independent of age, Townsend deprivation index, BMI and the first ten principal components of ancestry. Similarly, among all male patients from English recruitment centers with available COVID-19 test reporting, each SD increase in free androgen index increased risk of a positive test (OR=1.14, 95% CI: 1.02-1.28, *p*-value 0.027), as well as severe COVID-19 infection (OR=1.15, 95% CI:1.01-1.31, *p*-value 0.037). In further sensitivity analysis, effects were not mitigated by additional adjustment for prevalent Hypertension and type 2 Diabetes Mellitus (Supplementary Figure 13). We did not identify any significant associations for Testosterone or SHBG status (Figure 5D, Supplementary Figure 14).

## Discussion

There are two important observations in the COVID-19 pandemic: the higher prevalence of severe complications in male individuals and the relative immunity in children. Our study identifies a link between male sex hormone signaling and regulation of the SARS-CoV-2 receptor ACE2 and co-receptor TMPRSS2 providing a potential explanation for these observations. Our results demonstrate that inhibitors of 5 alpha reductases, which dampen androgen signaling, can reduce ACE2 levels and thereby decrease internalization of the viral spike-RBD. These drugs, commonly prescribed for prostatic disorders, have good safety profiles and show repurposing potential for the treatment of COVID-19.

The most lethal complication of COVID-19 is multiorgan failure affecting the lungs, the kidneys and the heart. Although there has been tremendous effort towards understanding the biology of SARS-CoV-2 infection at the molecular level, the use of relevant experimental models has been limited. Modeling and studying pathophysiology with cell types that are not directly affected by SARS-CoV-2 could potentially delay the identification of effective therapeutic targets. Taking advantage of directed hESC differentiation, we generated scalable cultures of disease-relevant human cardiac cells and performed high-throughput screening to identify drugs that regulate ACE2 expression in these cell types. These experiments underscore the potential of hESC-derived cardiac cells for in-depth investigation of cell-type specific processes and offer a framework for rapid identification and validation of therapeutically-relevant compounds. Other reports on the use of hESC-derived cell or organoid models highlight the utility of hESC-derived cells in modeling SARS-CoV-2 infection and COVID-19 pathophysiology^33^.

Results from our *in vitro* high-throughput screening of the FDA-approved drugs enable us to develop a novel deep learning strategy to screen millions of drug-like compounds *in silico* and identify candidates predicted to show superior potency and efficacy. The diverse pharmacokinetic properties in these candidates may enable the development of drugs with improved distribution to disease-relevant tissues. Further experiments are needed to validate these compounds and characterize their pharmacokinetic and pharmacodynamic properties *in vitro* and *in vivo*.

A common characteristic among our validated hit compounds is their ability to target androgen signaling. Analysis of disease outcomes in COVID-19 patients in two independent cohorts revealed a significant association between elevated free androgen and COVID-19 complications pointing to a possible link between androgen-mediated ACE2 regulation and disease severity. Pathway and gene target analysis on compounds that reduce ACE2 levels also highlighted the regulatory roles of peptidase pathways. Interestingly, protein interaction maps suggested a possible cross-talk between AR signaling pathways, inflammatory markers and peptidases relevant to the viral receptor and co-receptors, offering insights into alternative pathways involved in ACE2 regulation.

Our FDA drug screen data revealed that many commonly used medications modulate ACE2 levels and could affect disease severity in COVID-19 patients. Further studies evaluating the relationships between these drugs and disease outcomes will be necessary to assess potential clinical impact and the need to substitute medications that might pose a heightened risk for COVID-19 patients.

In conclusion, our results provide key insights into ACE2 regulatory mechanisms, present strong molecular and clinical evidence for the role of androgen signaling in SARS-CoV-2 infection and identify potential therapeutic candidates for the treatment of COVID-19.

## Methods

### Cell culture and differentiation

H9 human embryonic stem cells were maintained and differentiated as described previously^10,12^. Briefly, human embryonic stem cells (hESCs) were maintained in mTeSR medium. hESCs were re-plated 72 hours prior to initiating differentiation. The cardiac differentiation was started with a mesoderm induction cocktail comprised of with 1.5 μM CHIR99021 (CHIR, Stem-RD), 20 ng/mL BMP4 and 20 ng/mL Activin A in RPMI (Cellgro) supplemented with B27 minus insulin, 2 mM GlutaMAX, 1x NEAA and 1x Normocin (InvivoGen) for 3 days (RPMI+B27 w/o insulin). Next, cells were treated with 5 μM XAV939 from day 3-6 in RB27-INS. From day 6 onward, differentiation of cells was carried out in RPMI supplemented with B27, 2 mM GlutaMAX, 1x NEAA and 1x Normocin (RPMI+complete B27). The protocol is outlined in supplementary Figure 1B.

Human pulmonary alveolar epithelial cells were purchased from Sciencell (cat#3200) and maintained according to the manufacturer’s instructions.

### High-throughput screening and drug candidate selection

hESC-derived cardiac cells were replated at day 25 of differentiation in 384 well plates. 72 hours after re-plating, the cells were treated with compounds from an FDA-approved chemical library (Selleckchem). 24 hours after treatment, cells were fixed stained with ACE2 antibody. High-throughput imaging and quantification of fluorescence signals were carried out using the In Cell Analyzer 2000 (GE Healthcare, USA). ACE2 signal intensity was normalized to total cell number within each well as measured by DAPI staining. Z-score calculation was performed and normalized for each plate.

### Spike-RBD cell internalization assay

24 hours after replating into 96 well plates, hESC-derived cardiac cells were treated with either vehicle or drug candidates. After 24 hours, 0.5 μM/mL human Fc-tagged recombinant spike-RBD protein with (Sino Biological Inc. 40592-V02H) was added to the medium and incubated for 30 minutes. The cells were subsequently fixed, permeabilized and stained with antibodies against ACE2 and human Fc receptor. Image analysis was performed as described above.

### Immunofluorescence staining

Cells were washed with PBSx3 and fixed with 4% paraformaldehyde for 30 minutes at 4 °C. Non-specific antigen binding was blocked by treating the cells with PBS+0.5% BSA prior to adding the primary antibodies (ACE2 or TMPRSS2, Proteintech cat#21115-1-AP and Novus Biologicals cat# NBP1-20984, respectively). Cells were incubated with the primary antibody solution overnight at 4 °C then washed with PBSx3. Secondary antibodies and fluorophore conjugated anti human Fc antibody were incubated for 30 minutes at room temperature. Finally, cells were washed with PBSx3 and stained with DAPI for nuclear counterstaining.

### Virtual high-throughput screening dataset

In order to train vHTS platform, we first split the data into training (n=1048), validation (n=131), and test datasets (n=131) (Supplementary Figure 2A-C), and used the validation set to evaluate increasingly complex models. We used Morgan fingerprints to represent the chemical features of each compound in the screened FDA-approved library (CircularFingerprint function available in DeepChem with SMILES as input). We tested a random forest regressor (scikit-learn), which failed to perform adequately (validation R^2^ score ~0). We then used an XGBoost model (XGBRegressor) with the following parameters: colsample_bytree = 0.3, learning_rate = 0.1, max_depth = 5, alpha = 10, n_estimators = 10. While the model performance improved (correlation coefficient of 0.1), it was largely driven by positive *z*-score values and the prediction of the negative side was substantially less reliable. We then tested graph convolutional neural networks after featurizing the molecules using ConvMolFeaturizer (DeepChem) and testing a default model architecture with dropout set to 0.2. This initial model was promising with an R of 0.05; therefore, we performed a systematic hyperparameter tuning using a grad search on the size of the graph convolutional and dense layers, as well as dropout rates. The best-performing model contained two graph convolutional (size 64) and dropout of 0.5 (and uncertainty set to true) and achieved an R of 0.1. We also tested message-passing models (MPNN) in a similar fashion; however, the performance was not improved. Therefore to further boost the performance of the models, we first focused on improving the dataset itself. In addition to the ACE2 *in vitro* screening data, we included data from five additional screens with the same library, added an inverse normal transformation when appropriate, and performed a multi-task learning. These modifications allowed the model to generalize better and learn faster with better R scores (this allowed us to reduce the dropout to 0.25). Second, instead of using a single model, we utilized a bagging ensemble model instead, where five models were trained on random samplings (with replacement) of training data. This ensemble model achieved an R of >0.2 on the validation dataset. We then continued training while the Pearson coefficient for validation set remained above 0.2 (early stopping). Finally, we tested this final ensemble model on the test dataset, which achieved a similar R=0.22 (*p*-value 0.01).

#### In silico screening

The final model described above was used to evaluate 9.2 million compounds from the ZINC15 database. We downloaded SMILES from this database and featurized them (same as above). The resulting predictions were variance normalized and the molecules with *z*-scores below −4 for ACE2 expression were selected as hits. This resulted in 169 potential small molecules.

#### Data visualization

To visualize the relationship between the identified hits in a lower dimension, we used Morgan fingerprints to represent all compounds and used UMAPs to project each compound down to 2 dimensions. For the screened ZINC15 database, we sampled 1 from 1000 compounds for visualization purposes. We also used the Morgan fingerprints to cluster *in silico* and *in vitro* hits using K-means clustering (number of clusters set to 15).

### Predicting drug-protein-pathway interactions

#### Identification of drug targets in FDA-approved library

Isomeric SMILES for each drug in the library were acquired from Selleckchem and used to run a similarity ensemble approach (SEA) library search. The SEA predicted targets were filtered, selecting human targets and predicted interaction *p*-values <0.05, which yielded 2150 predicted proteins targeted by the drug library.

#### Target drug selection

The normalized *z*-score values reported for all the compounds were first transformed to N(0,1) using the bestNormalize package (v1.4.0) in R (v3.5.1). The treatments with transformed *z*-scores smaller than −1.5 were selected, which resulted in 41 compounds.

#### Target gene selection

For every compound, possible target genes were identified as above. Weighted combined z-scores were then calculated for each gene by combining normalized *z*-scores across all treatments. The *p*-values were then calculated based on the combined z-scores and adjusted using p.adjust (method=FDR). As an orthogonal approach, for each gene, we recorded the number of treatments with negative normalized *z*-scores as well as the total number of compounds predicted to target that gene. Using the sum of counts for all other genes and drugs, we performed a Fisher’s exact test to evaluate the degree to which negative *z*-scores were enrichment among the treatments likely to affect a gene of interest. As expected, the two *p*-values, i.e. combined *z*-score and Fisher’s, are generally correlated (R=0.6, *p*<1e-200).

#### Gene-set enrichment analysis

We used the combined *z*-scores across all genes to identify pathways and gene-sets that are associated with changes in ACE2 expression. For this analysis, we used our iPAGE toolkit^34^, in conjunction with annotations from MSigDB and Gene Ontology (GO). The following parameters were set: --ebins=9 --nodups=1 --independence=0.

#### Drug-protein-pathway analysis

For drugs with a normalized z-score ≥-1, drug-protein interactions per biological pathway were analyzed. The SEA predicted drug-protein interaction dataset was filtered to exclude drugs with a normalized z-score ~-1. Protein pathway gene lists were generated per gene set, comprised of the gene set genes present in the pre-ranked gene list. For each protein pathway gene list, drug-protein matrices were plotted by the SEA *p*-value for the interaction. The reported interaction significance scores represent the.

### Single cell sequencing data analysis

#### Data sources

Single cell transcriptomics data and associated metadata were obtained from Tissue Stability Cell Atlas (https://www.tissuestabilitycellatlas.org/), Gene Expression Omnibus (GEO; https://www.ncbi.nlm.nih.gov/), and Single Cell Portal (https://singlecell.broadinstitute.org/single_cell). Human lung and esophageal datasets containing samples from 5 and 6 previously healthy, post-mortem donors, respectively, were obtained from research published by Madissoon et al.^14^ A human heart dataset containing samples from 12 previously healthy, post-mortem donors was obtained from research published by Wang et al, GSE109816^15^. A human colon epithelia dataset containing biopsy samples from 12 healthy individuals were obtained from research published by Smillie et. al.^13^

#### Quality control

Cells were identified as poor quality and subsequently removed based on the criteria implemented by the original authors of each dataset.

##### Lung and esophagus datasets

RData objects deposited to Tissue Stability Cell Atlas were pre-filtered by the authors. Filtering criteria can be found in the methods of Madissoon et al.^14^ Briefly, cells were excluded if they did not meet all the following criteria: more than 300 and fewer than 5,000 genes detected, fewer than 20,000 UMI, and less than 10% mitochondrial reads. Genes were removed if they were detected in fewer than three cells per tissue.

##### Colon dataset

The gene by cell counts matrix deposited to GEO was pre-filtered by the authors. Filtering criteria can be found in the methods of Wang et al.^15^ Briefly, cells were removed if they did not meet all the following criteria: detected at least 500 genes, UMI’s within 2 standard deviations of the mean of log10UMI of all cells, unique read alignment rate at least 50%, and fewer than 72% mitochondrial reads. Further, cardiomyocytes (CM) from CM-enriched datasets were included if UMIs detected were greater than 10,000.

##### Heart dataset

Based on the methods of Smillie et al,^13^ cells were removed if they did not meet one of the following criteria: minimal expression of 500 genes per cell; nUMIs within two standard deviations from the mean of log_10_ nUMIs of all cells; unique read alignment rate (# of assigned reads/ # total aligned reads) greater that 50%; and a mitochondrial read percentage of less than 72%.Mitochondrial genes were subsequently removed from the dataset prior to dimensionality reduction.

#### Data integration, dimensionality reduction and cell clustering

Different methods available in Seurat^35^ were implemented respective to individual datasets.

##### Lung and esophagus datasets

Batch correction, data normalization variable gene identification, data scaling, principal component analysis (PCA) and uniform manifold approximation and projection (UMAP) dimensionality reduction was performed by the original authors and included in the RData objects deposited to Tissue Stability Cell Atlas. The original UMAP coordinates generated by the authors were used for any visualizations.

##### Colon Dataset

Batch correction by patient sample was performed using the Seurat v3 integration functions. The dataset was split by “Subject”, individual gene matrices were log normalized using a scaling factor of 10,000 and 2,000 variable features were identified per individual gene matrix using variance stabilizing transformation. 2,000 integration anchors were found using the first 30 dimensions of the canonical correlation analysis and individual datasets were integrated using the same number of dimensions. PCA was performed and UMAP coordinates were found based on the first 20 principal components.

##### Heart Dataset

Batch correction by patient sample was performed using mutual nearest neighbor (MNN)^36^ matching through the “RunFastMNN” function from the SeuratWrappers package. The dataset was log normalized using a scaling factor of 10,000 and 2,000 variable features were identified per individual gene matrix using variance stabilizing transformation. “RunFastMNN” was performed on the dataset split by “Individual” and UMAP coordinates were found based on the first 30 MNN dimensions. A shared nearest neighbor graph was constructed with the same number of MNN dimensions and clusters were identified using clustering resolution of 0.2.

#### Cell type identification and gene expression analysis

##### Lung, esophagus and colon datasets

Cell type cluster annotations determined by the original authors were available through the metadata downloaded with each dataset. These original cluster annotations were used for any visualizations.

##### Heart Dataset

Clusters were annotated based on cluster specific expression of marker genes identified in the original publication. Cluster markers were identified using a Wilcoxon Rank Sum test.

Following cell type annotation, gene dropout values were imputed using adaptively-thresholded low rank approximation (ALRA)^37^. The rank-k approximation was automatically chosen for each dataset and all other parameters were set as the default values. The imputed gene expression in shown in all plots and used in all downstream analysis.

#### Gene signature scoring

The Seurat “AddModuleScore” function was used to score each cell in the datasets for their expression of high confidence drug target genes (Figure 2C) and genes associated with AR signaling (Supplementary Table 3). 100 control genes selected from the same bin per analyzed gene were used to calculate each module score. “Upstream AR Activators” were identified through literature search^24–27^, “AR Signaling” genes were taken from the androgen receptor signaling pathway gene ontology term (GO:0030521), and “Common AR Target Genes” are the common “core” target genes transcribed by AR activated identified by Jin et. al.^28^ through comparison of multiple microarray studies.

### Bulk RNA sequencing analysis

The bulk RNA sequencing dataset analyzed here was previously reported^12^. Briefly, human pluripotent stem cell (H9 with knock-in MYH6:mCherry reporter)-derived cardiomyocytes were prepared as described previously^10,12^. Reporter tagged cardiomyocytes were isolated from the negative fraction of the culture by FACS sorting. Both the purified cardiomyocytes and negative fractions were prepared and sent for bulk RNA sequencing. The resulting datasets included two negative fraction biological replicates and two positive fraction biological replicates. Gene expression between the cardiomyocytes and non-cardiomyocytes was compared by averaging the read count per gene and normalizing by the average read count of GAPDH.

### Protein-protein interaction network construction

Protein-protein interaction network analysis was performed using the Search Tool for the Retrieval of Interacting Genes (STRING) database. The minimum required interaction score was set to 0.7 corresponding to high confidence with the edge thickness indicating the degree of data support from the following active interaction sources: textmining, experiments, databases, co-expression, neighborhood, gene fusion and co-occurrence.

### Signaling pathway illustration

SEA predicted drug-protein interactions were connected by lines. Lines were dashed if the MaxTC score was below 1. Proteins corresponding with the 30 significantly enriched genes from 2C were highlighted in yellow. To identify pathways involving our candidate proteins, we combined data from the Kyoto Encyclopedia of Genes and Genomes (KEGG)^38^ database and previous reports in the literature. Adobe Illustrator 24.1 was used for visualization.

### Transcriptional changes associated with AR down-regulation

We used an existing AR ChIP-seq dataset generated in LNCaP cells to identify direct transcriptional targets of AR. We downloaded processed peaks from the Gene Expression Omnibus (GSM3148987) and lifted the peaks from hg19 over to hg38. We then used annotatePeak function (package ChIPseeker v1.8) to identify peaks 5kb downstream or upstream of transcription start sites (TSS). The genes with AR binding to this 10kb window around their TSS were selected as direct AR-bound targets.

We also used an RNA-seq dataset generated using RNA-mediated knockdown of AR (SRR7120653, SRR7120649, SRR7120646, and SRR7120642). We downloaded raw fastq files and aligned them to the transcriptome (gencode.v28) using Salmon (v0.14.1 using --validateMappings and −l ISR flags). We then used DESeq2 (1.22.2) to compare gene expression changes in response to AR knockdown. We then used the AR-bound gene-set from the previous step to assess how the AR targets respond to its down-regulation. For this, we used our iPAGE tool^34^, which uses mutual information (MI) and an associated *z*-score to assess the enrichment/depletion patterns of a gene-set across gene expression modulations. To visualize this data, we included a volcano plat, with genes of interest shown in red. For this analysis, we divided log-fold changes into nine equally populated bins, which were also included along with the patterns of enrichment and depletion across the data.

### Yale New Haven Hospital patient dataset

EPIC’s SlicerDicer feature was used to collect the Yale New Haven Hospital (YNHH) electronic health record anonymous aggregate-level data. Using this feature, a total of 8657 patients had a COVID-19 ICD-10 diagnosis code at YNHH during November 28, 2019-April 28 2020 time period. Among them, 1577 patients had a Troponin T checked during the same encounter and were selected for subsequent analysis. Out of 1577 patients, 787 were female and 790 were male.

Next, we focused on males above age 40 for subsequent risk factor analysis. Risk factors associated with disease outcome, including age, BMI, status of hypertensive disorder, status of diabetes mellitus disorder and prostate disease were collected as dichotomized variables during data download. Variable cutoffs were set as follows: age, 40-65y vs. 65+ y, BMI, <30 vs. >=30, troponin T <0.01 ng/ml vs. >=0.01 ng/ml. Diabetes, hypertension and prostatic disease were binary originally and were collected as a visit diagnosis. Appropriate SNOMED-CT (systematized nomenclature of medicine–clinical terms) were used to include or exclude patients according to their pre-existing conditions. The following terms were used to identify patients with pre-existing conditions in the database: “disorders of glucose metabolism” to select the patients with any form of the Diabetes Mellitus, “Hypertensive disorder” to select the patients with any form of hypertension and “hyperplasia of prostate or Neoplasm of Prostate” to select patients with disorders of prostate gland. The study was in accordance with Institutional Review Board policy.

A total of 681 subjects exist in this dataset with 239 high troponin and 442 low troponin cases. The interdependence of all variables was explored through their correlation matrix (Supplementary Figure 10B) and pairwise Fisher’s exact tests. To find the optimal model, we used a backward selection approach. We started with a logistic regression model with troponin as response and all the other variables and their pairwise interaction terms as covariates. In successive steps, least significant terms were eliminated one at a time and the model was refitted until all remaining terms were significant at p<0.05. Odds of having abnormal troponin T in the baseline group (40-65y, no prostate disease, low BMI, no diabetes, no hypertension, n=58) was 0.038.

### UK Biobank dataset

Individual-level longitudinal phenotypic data from the UK Biobank, a large-scale population-based cohort with genotype and phenotype data in approximately 500,000 volunteer participants recruited from 2006-2010 was used^39^. Baseline assessments were conducted at 22 assessment centers across the UK using touch screen questionnaire, computer assisted verbal interview, physical tests, and sample collection including for DNA. Additional details regarding the study protocol are described online (www.ukbiobank.ac.uk).

Of 488,377 individuals genotyped in the UK Biobank, we used data for 311,857 participants from English recruiting centers where COVID-19 testing was reported in the UK Biobank, as done previously^40^. Additionally, we further filtered to individuals with white British ancestry consenting to genetic analyses, with genotypic-phenotypic sex concordance, without sex aneuploidy, and removed one individual from each pair of 1^st^ or 2^nd^ degree relatives selected randomly. Secondary use of the data was approved by the Massachusetts General Hospital institutional review board (protocol 2013P001840) and facilitated through UK Biobank Application 7089.

#### UK Biobank androgen indices and phenotypic covariates

Free androgen index was calculated in the UK Biobank by using total testosterone and sex hormone-binding globulin (SHBG) as in the equation: Free androgen index= 100*(total Testoterone/SHBG). Testosterone (nmol/L), SHBG (nmol/L), and free androgen index (unitless) phenotypes were observed to have varying distributions by sex, with outliers. Sex-specific extreme outliers for each phenotype were identified and excluded by adjusting the traditional box and whisker upper and lower bounds, accounting for skewness in the phenotypic data identified using the Robustbase package in R (setting range=3) (https://cran.r-project.org/web/packages/robustbase/robustbase.pdf). After excluding extreme outliers, we performed inverse rank normalization of each phenotype by sex to a mean of 0 and SD 1 for use in analysis (Supplementary Figure 11B).

Definitions for phenotypic covariates included in sensitivity analysis are included in Supplementary Table 4. In brief, hypertension was defined by self-reported hypertension and billing codes for essential hypertension, hypertensive disease with and without heart failure, hypertensive heart and renal diseases, and secondary hypertension. Type 2 diabetes was defined by billing codes for non-insulin-dependent diabetes mellitus or self-reported type 2 diabetes. Enlarged prostate was defined by combining self-reported enlarged prostate and billing codes for prostatic hyperplasia.

#### UK Biobank COVID-19 case/control models

COVID-19 phenotypes used in the present analysis are from UK Biobank downloaded on May 2, 2020 and include combined nose/throat swabs and lower respiratory samples analyzed using PCR. Four case/control definitions were used in analysis to iterate over COVID-19 susceptibility and severity across both tested individuals and all individuals with reported COVID-19 testing from the UK Biobank English recruitment centers^40^. COVID+ cases were defined as any individual with at least one positive test. Severe COVID+ cases were defined as any individual with at least one positive test who also had evidence from their microbiological record that they were hospitalized inpatient. Additionally, individuals with COVID-19 of unknown severity were excluded from the Severe COVID+ analyses. Thus, the four definitions were as follows: for individuals tested with COVID: comparing 1) COVID+ vs. COVID-test results, 2) Severe COVID+ vs. COVID-; and for all white British individuals from the UK Biobank English recruitment centers with available COVID-test reporting: 3) COVID+ vs. All, 4) Severe COVID+ vs. All.

#### UK Biobank statistical models

Association of the three androgen indices (Free androgen index, testosterone, and SHBG) with COVID-19 across the four case/control definitions was performed using logistic regression in R-3.5. Three main association models were used for analysis: 1) an unadjusted model, 2) a sparsely adjusted model using age, normalized Townsend deprivation index (a marker of socioeconomic status), and the first ten principal components of genetic ancestry, and 3) a fully adjusted model that additionally adjusted for normalized body mass index (BMI). Analyses were performed across males and females as well as separately by sex. Several additional sensitivity analyses were performed to assess for the robustness of associations. In particular, analyses were performed additionally adjusting for prevalent hypertension and prevalent type 2 diabetes, showing consistent associations.

### Statistical methods

Experiments were carried out in a minimum of 3 biological replicates unless specified otherwise. Two-tailed student’s t-test was performed to compare treatment groups unless otherwise indicated in the methods section or legends. Error bars in s represent standard error of the mean.

## Acknowledgements

We are grateful to the UCSF program for breakthrough biomedical research and Sandler foundation for the grant to F.F. that supported this work. F.F. is also supported by the NIH Director’s New Innovator Award (DP2NS116769) and the National Institute of Diabetes and Digestive and Kidney Diseases (R01DK121169). H.M. is supported by Larry L. Hillblom Foundation postdoctoral fellowship. S.M.Z. is supported by the National Institutes of Health’s NHLBI under award number 1F30HL149180-01 and the National Institutes of Health’s Medical Scientist Training Program at the Yale School of Medicine. H.A. is supported by the NIH postdoctoral fellowship F32GM133118 and the postdoctoral grant from the program for breakthrough biomedical research grant and the Sandler foundation at UCSF. H.G. is supported by the National Cancer Institute (R01 CA240984) and the National Institute of General Medical Sciences (R01 GM123977). P.N. is supported by a Hassenfeld Scholar Award from the Massachusetts General Hospital, and grants from the National Heart, Lung, and Blood Institute (R01HL1427, R01HL148565, and R01HL148050) and Fondation Leducq (TNE-18CVD04).

## Author contributions

Z.G., H.M, M.R., R.S., S.F., A.K., J.R. performed the experiments, H.A. performed statistical analysis S.M.Z, H.Z., P.N. performed the UKBB study analysis, H.G. performed pathway analysis and in silico library screen, F.F.: designed, conceptualized, performed and supervised the experiments, Z.G., H.M, M.R., R.S. H.G., F.F.: wrote the manuscript.

## Data availability

The supporting raw data of this study are available from the corresponding authors upon request.

---

**Supplementary Figure 1.**
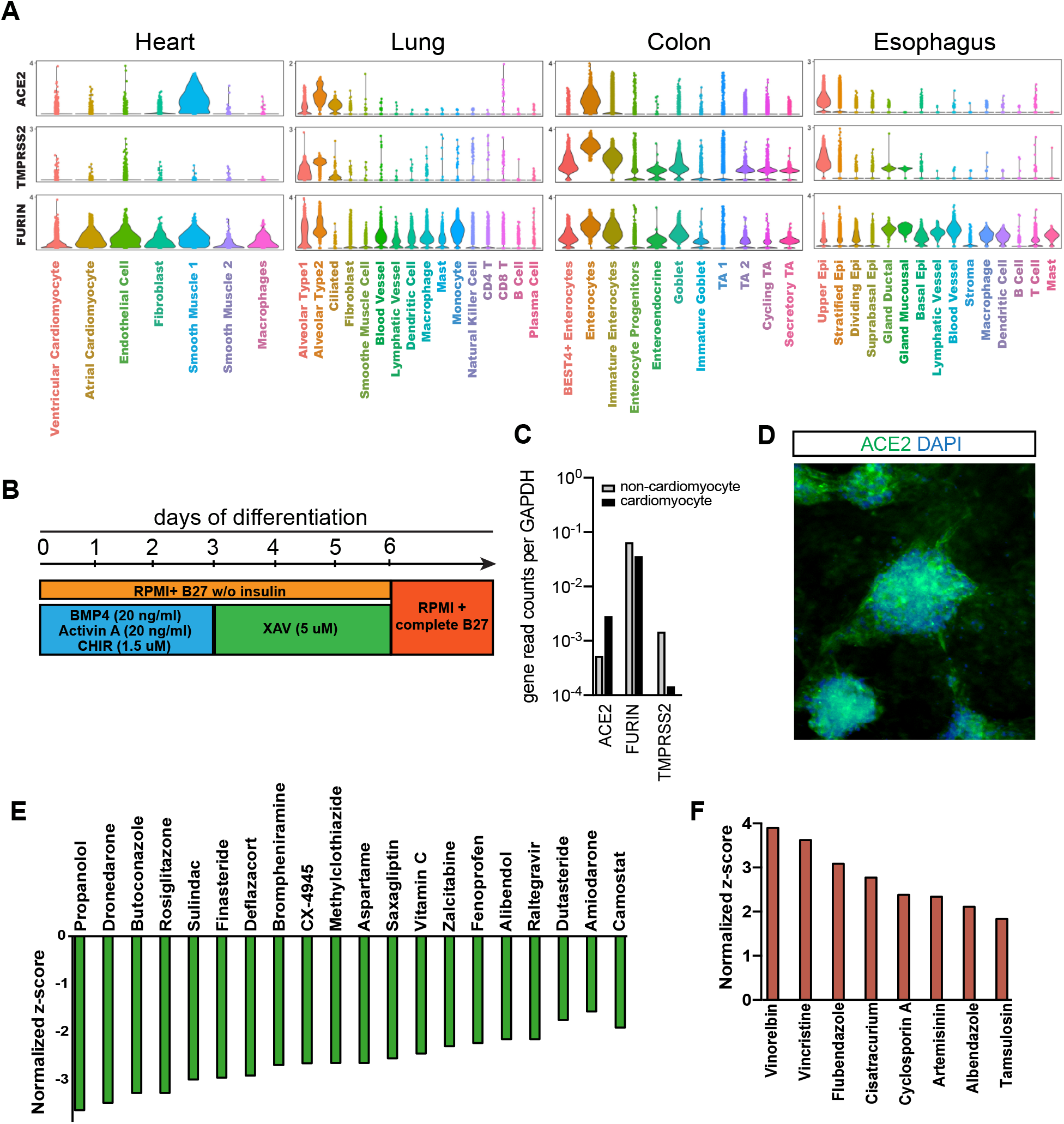
Cell type specific Expression of SARS-CoV-2 receptors. **A)** Violin plots showing the cell type specific expression of ACE2, TMPRSS2 and FURIN for human adult heart, lung colon and esophagus. The expression is displayed as the log normalized values. B) Schematic representation of cardiac differentiation protocol **C)** Expression of ACE2, FURIN, and TMPRSS2 in hPSC-derived cardiomyocytes and non-cardiomyocytes. Expression is normalized by GAPDH read count. **D)** Immunofluorescence imaging of cardiac cells stained with ACE2 antibody. **E-F)** Normalized z-score of compounds in FDA-approved library that up/down-regulate ACE2 in human cardiac cells.

**Supplementary Figure 2.**
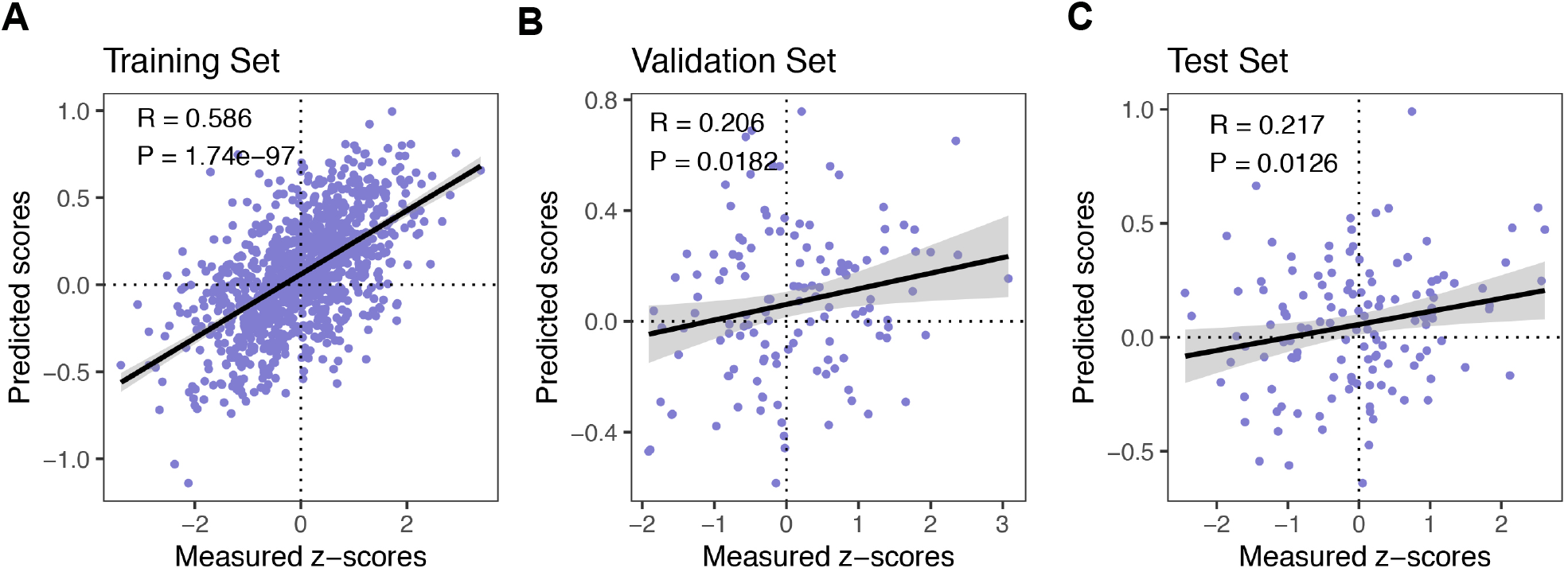
Assessment of the final ensemble model of GCNNs. Comparison of predicted and measured *z*-scores for the training (**A**), validation (**B**) and test (**C**) datasets in virtual high-throughput screening experiment. Also shown are the Pearson correlation coefficients and their associated *p*-values.

**Supplementary Figure 3.**
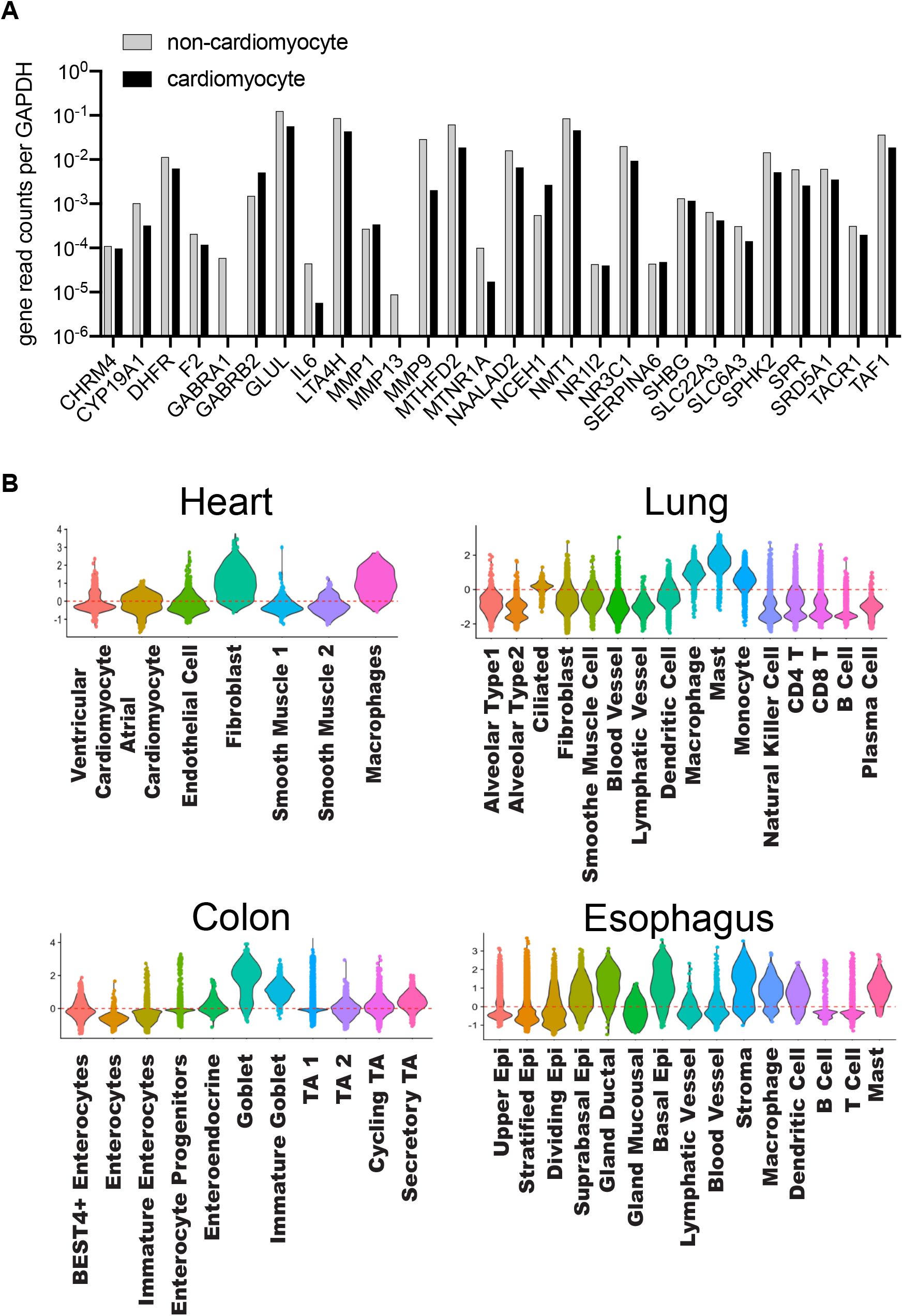
Cell type specific expression of predicted targets. **A)** Expression of predicted targets in hPSC-derived cardiomyocytes and non-cardiomyocytes. Expression is normalized by GAPDH read count. **B)** Violin plots showing the cell type specific module scores for the 30 predicted gene targets for human adult heart, lung colon and esophagus.

**Supplementary Figure 4.**
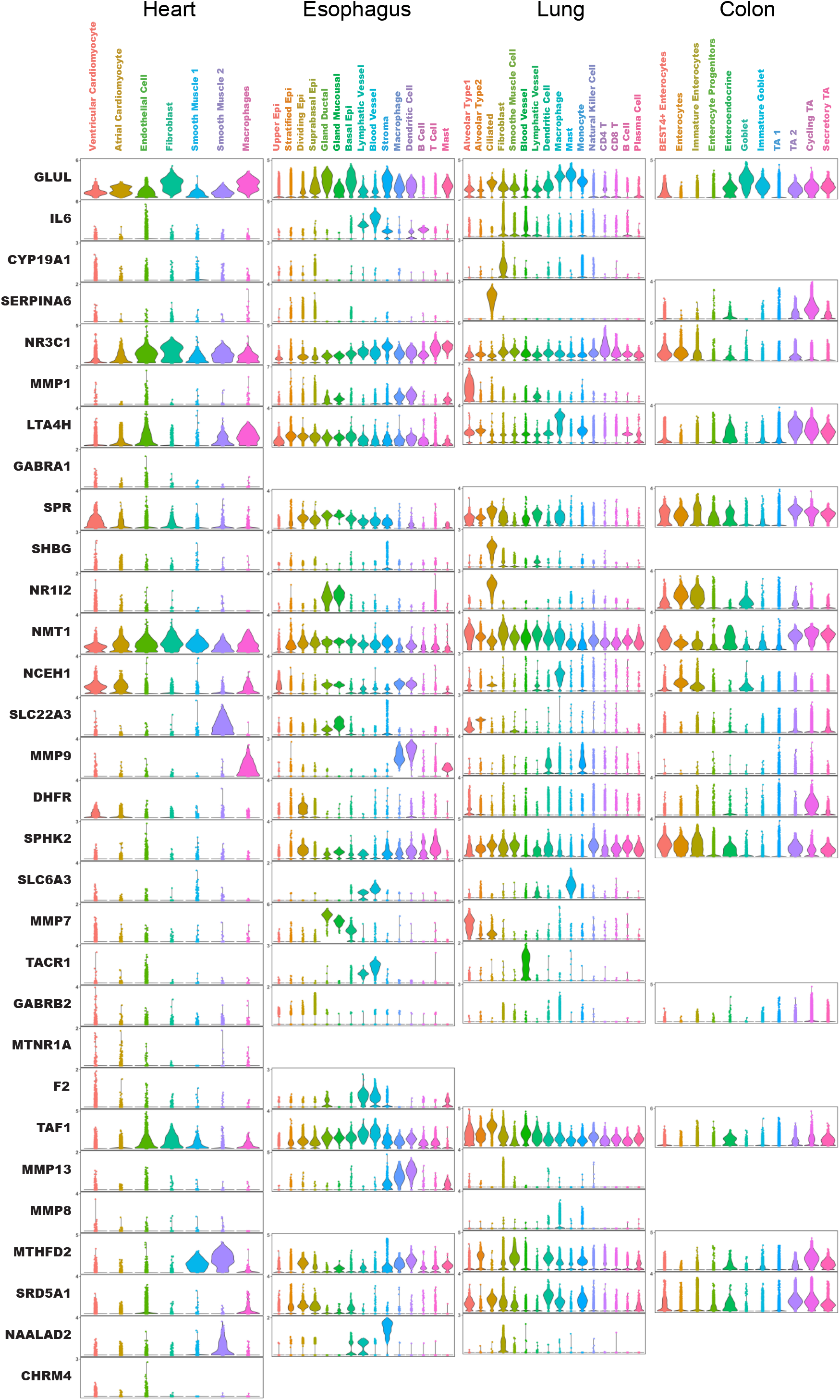
Cell type specific expression of individual predicted target genes. Violin plots showing the cell type specific expression of the 30 predicted gene targets for each organ. The expression is displayed as the log normalized values.

**Supplementary Figure 5.**
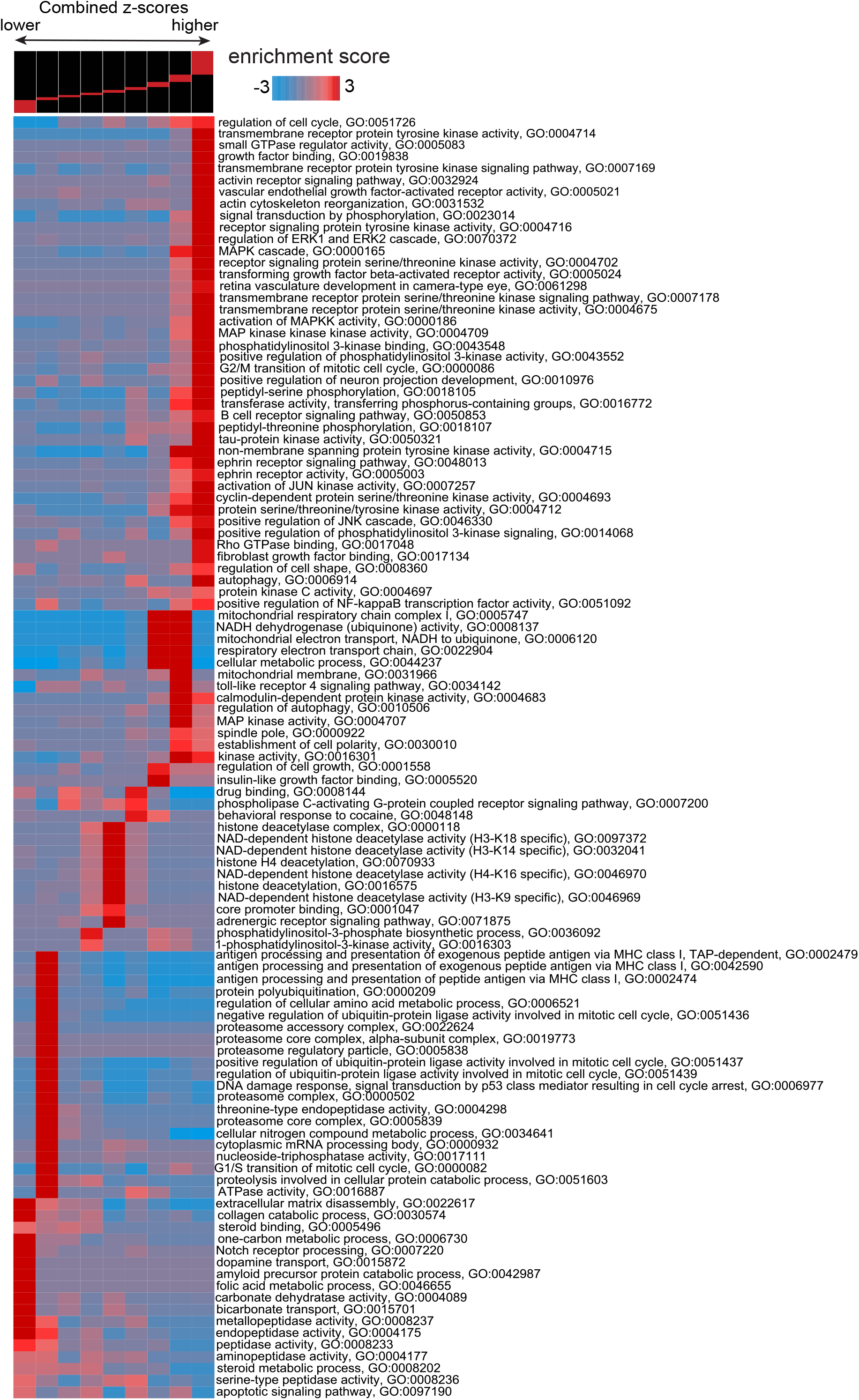
Pathway analysis of target proteins. Pathways discovered by iPAGE and their pattern of representation across proteins associated with high/low z-scores in the FDA-approved library screen. Each expression bin includes genes within a specific range of expression values (top panel). Bins to the left contain genes associated with lower z-scores and the ones to the right contain genes associated with higher z-scores. Rows correspond to pathways and columns to expression bins. Red entries indicate enrichment of pathway genes in the corresponding expression bin.

**Supplementary Figure 6.**
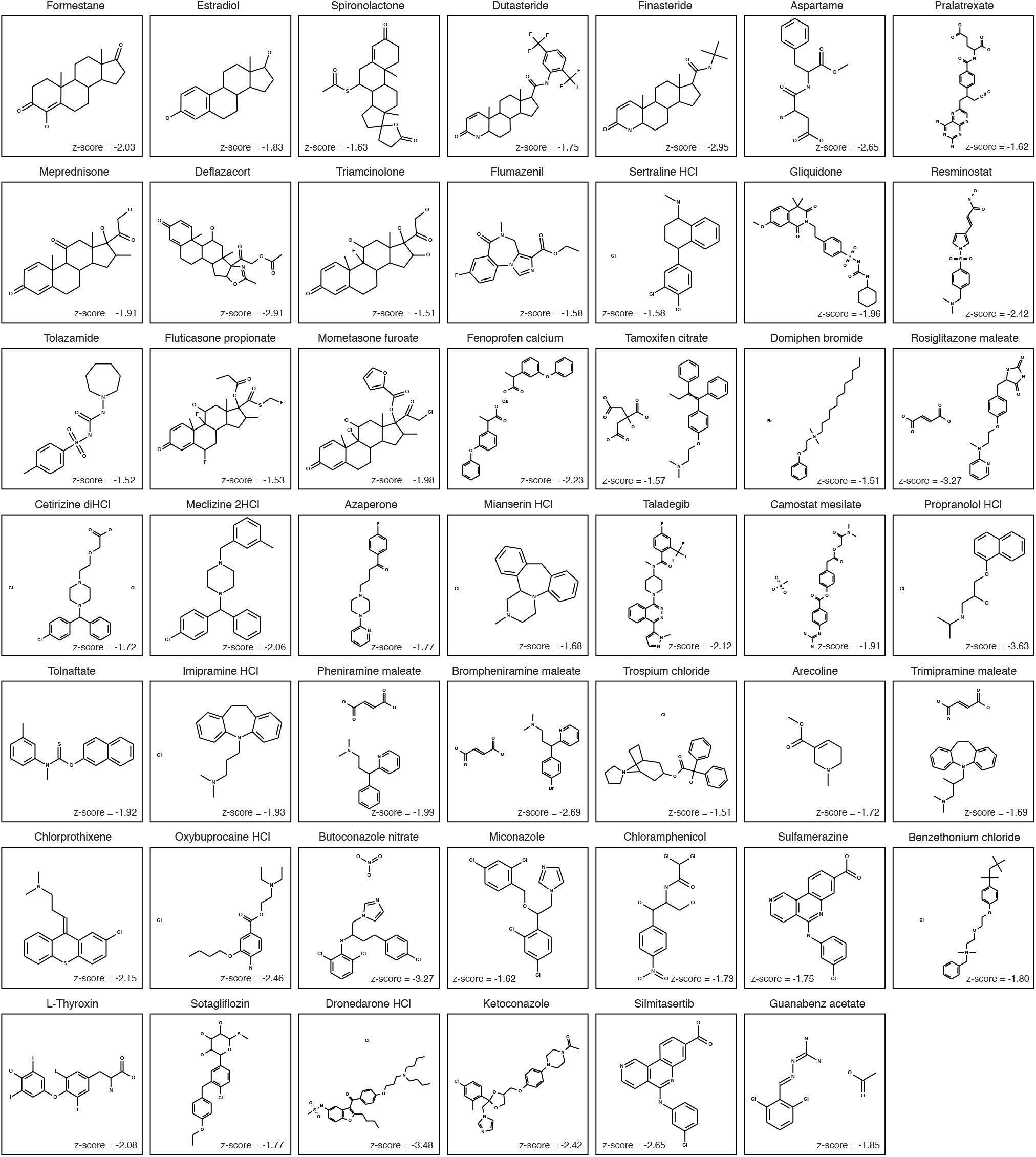
Chemical structure of hit compounds that interact with predicted targets. Canonical SMILES and Punchem sketecher were used to draw the structures.

**Supplementary Figure 7.**
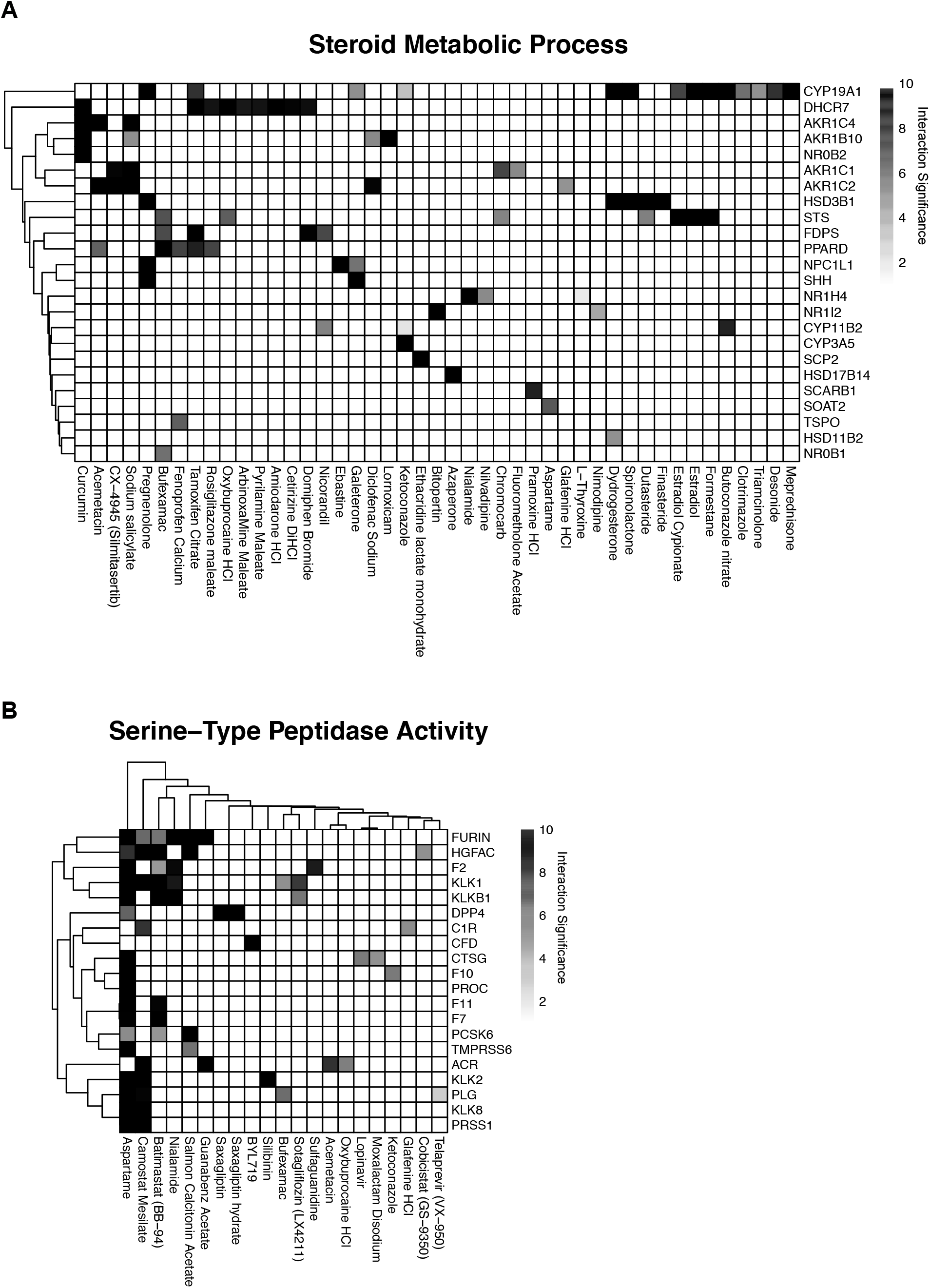
Drug-target interactions in selected GO terms. **A)** Drug-protein interaction matrix for GO:0008202 steroid metabolic process gene set. **B)** Drug-protein interaction matrix for GO:0008236 serine-type peptidase activity gene set.

**Supplementary Figure 8.**
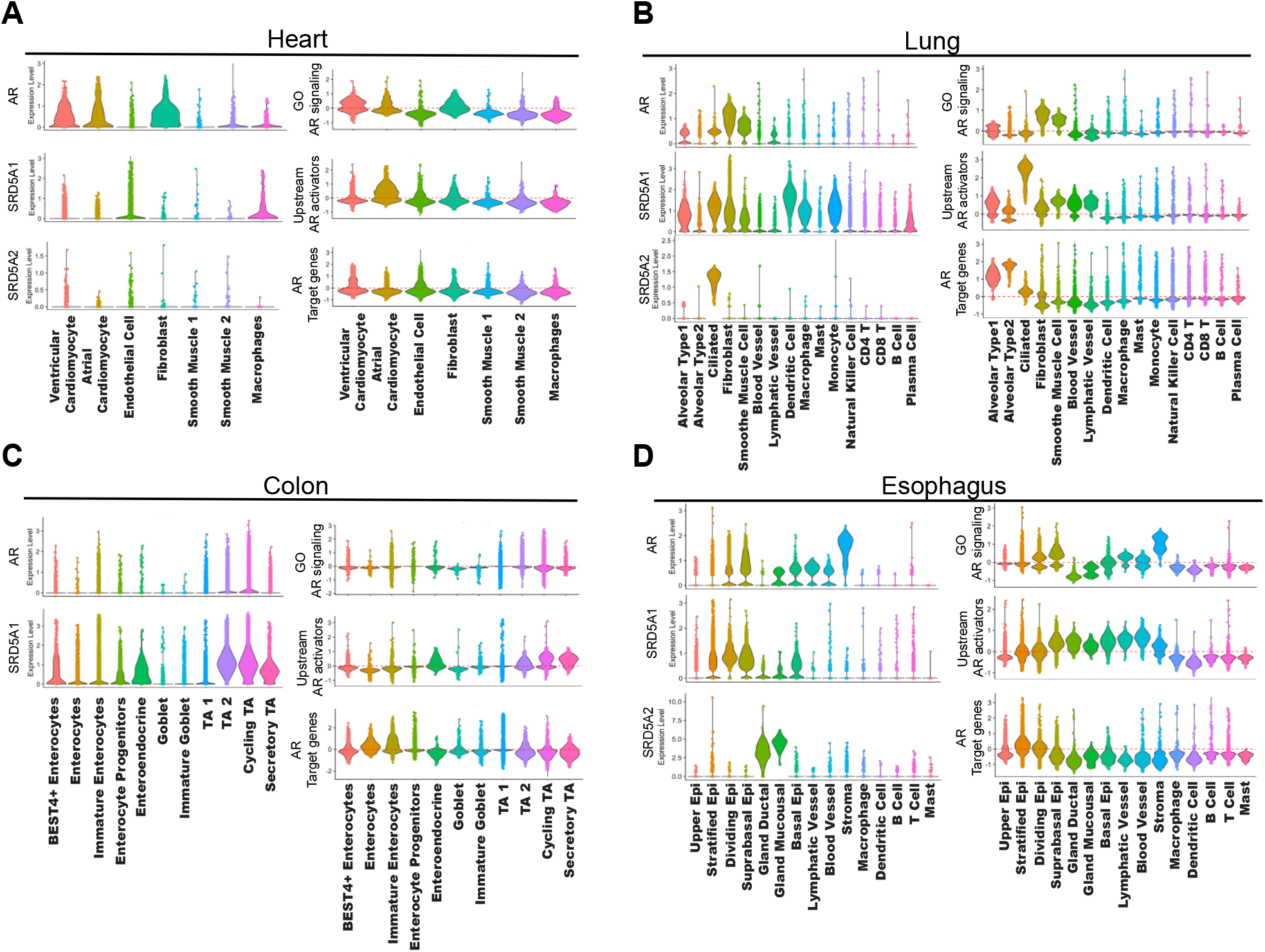
Cell type specific expression of AR signaling modulators. **A)** Violin plots showing the cell type specific expression of AR, SRD5A1, SRD5A2 and cell type specific module scores for 8 RTK’s upstream of AR activity, 30 genes associated with the androgen receptor signaling pathway and 34 common downstream targets of AR transcriptional regulatory activity in the human adult heart. **B)** Violin plots showing the cell type specific expression of AR, SRD5A1, SRD5A2 and cell type specific module scores for 8 RTK’s upstream of AR activity, 30 genes associated with the androgen receptor signaling pathway and 34 common downstream targets of AR transcriptional regulatory activity in the human adult lung. **C)** Violin plots showing the cell type specific expression of AR, SRD5A1, SRD5A2 and cell type specific module scores for 8 RTK’s upstream of AR activity, 30 genes associated with the androgen receptor signaling pathway and 34 common downstream targets of AR transcriptional regulatory activity in the human adult colon. **D)** Violin plots showing the cell type specific expression of AR, SRD5A1, SRD5A2 and cell type specific module scores for 8 RTK’s upstream of AR activity, 30 genes associated with the androgen receptor signaling pathway and 34 common downstream targets of AR transcriptional regulatory activity in the human adult esophagus.

**Supplementary Figure 9.**
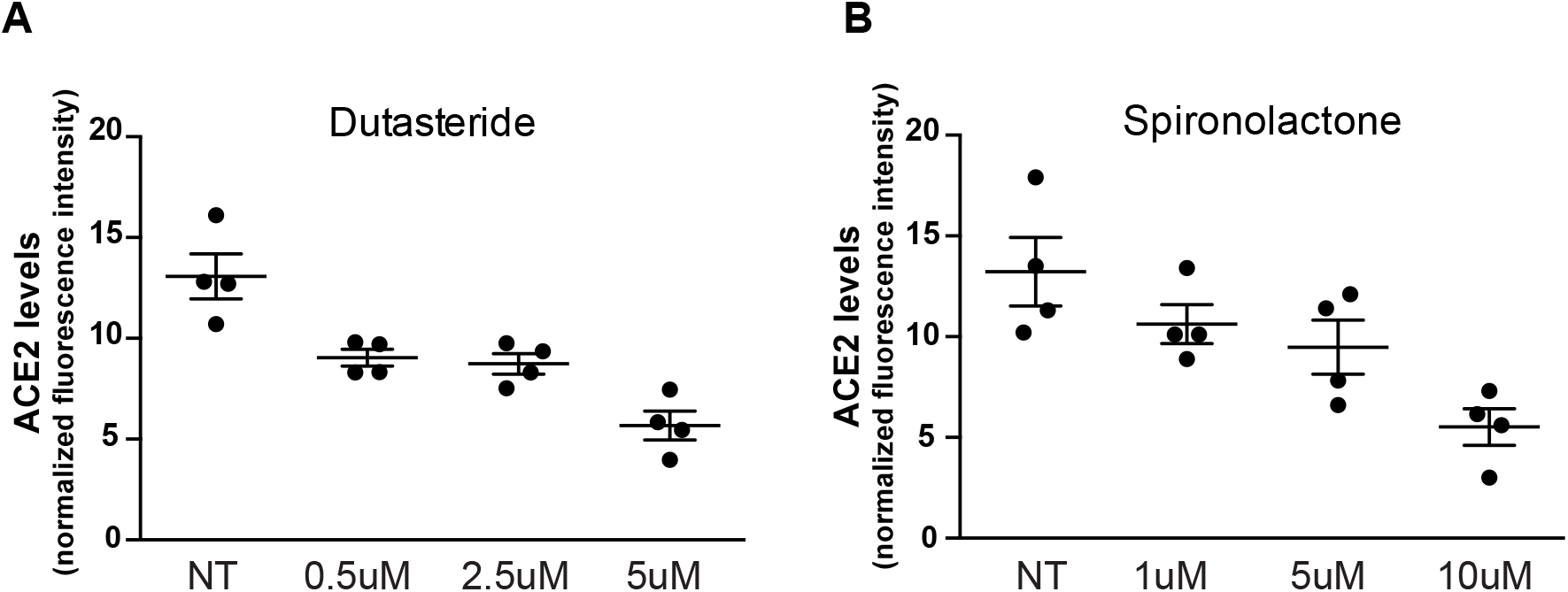
Dose specific effect of selected AR modulators on ACE2 levels. Dose response analysis for dutasteride (**A**) and spironolactone (**B**)

**Supplementary Figure 10.**
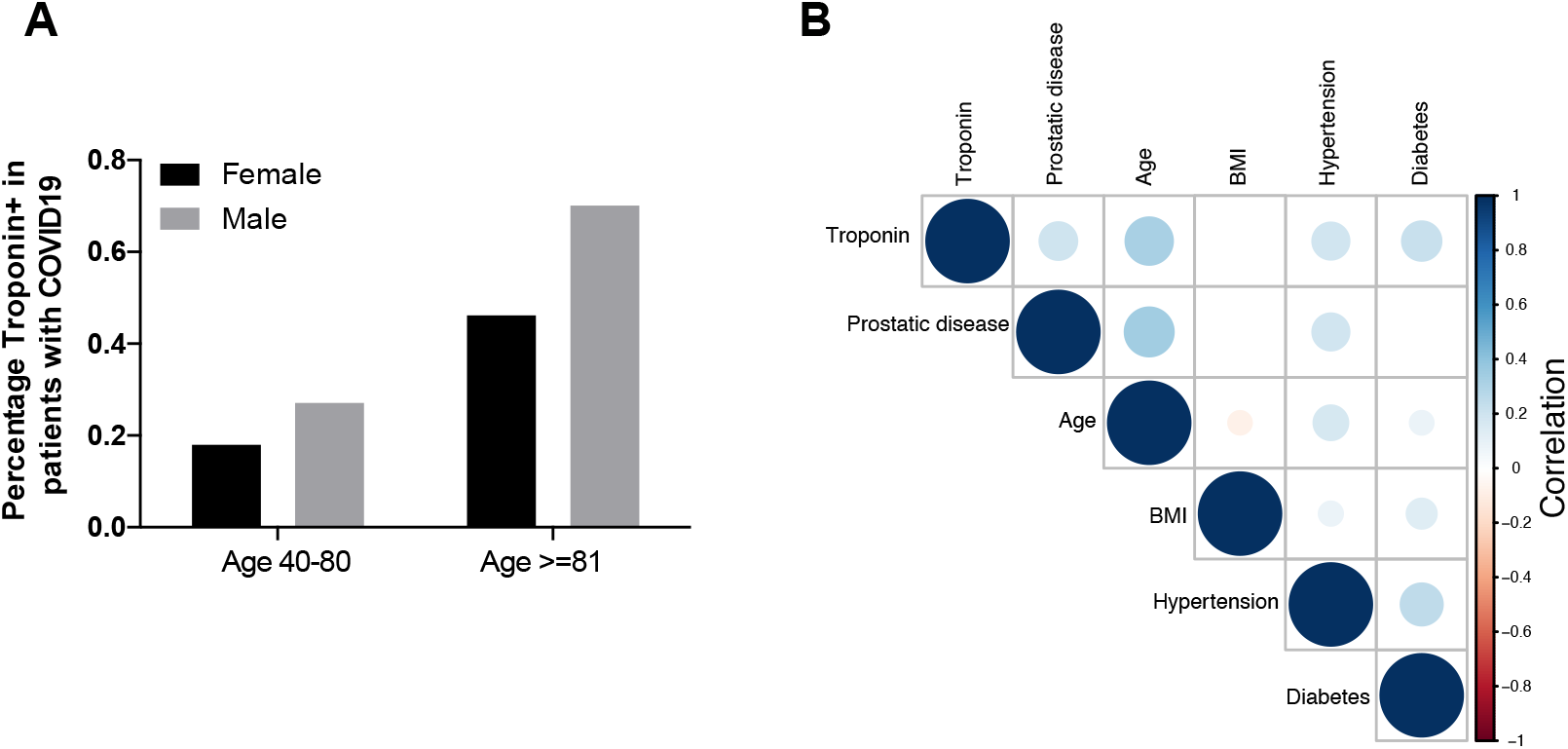
Effects of risk factors on abnormal troponin T in COVID-19 patients. **A.** Distribution of sex among patients with abnormal troponin T. **B.** Correlogram of variables in the Yale Covid-19 patients dataset. The size and color of each circle illustrates the direction and strength of the correlation. Only significant correlations (at p<0.05) are painted.

**Supplementary Figure 11.**
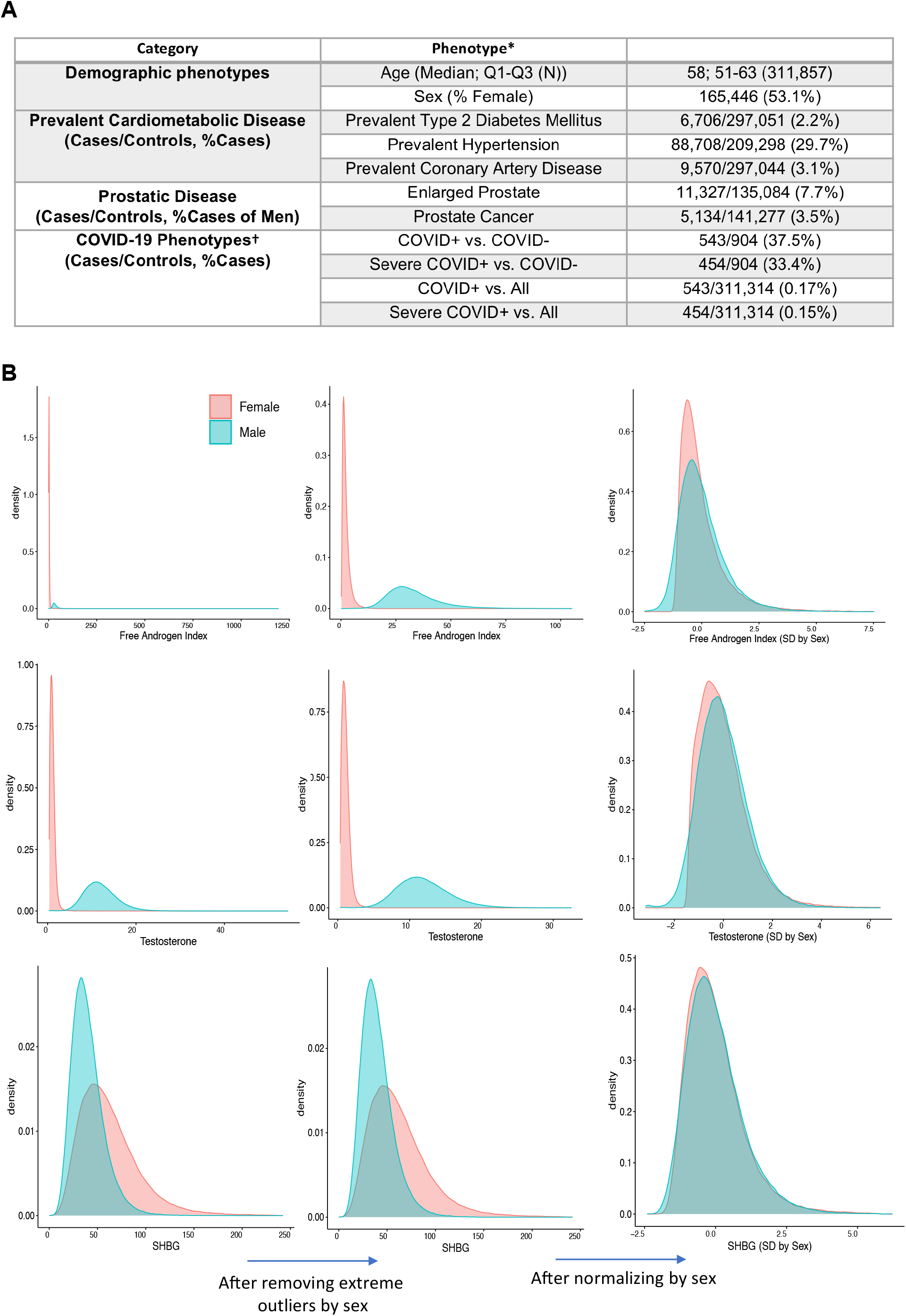
Distribution of demographic and androgen indices in UKBB cohort. **A.** Baseline characteristics among analyzed UKBB participants. * These values reflect the samples used in analysis after quality control filters were applied. †COVID+= Tested positive for Coronavirus disease 2019, COVID-=Tested negative for Coronavirus disease 2019, all= all individuals with reported Covid-19 testing from the UKBB English recruitment centers. **B.** Distribution of androgen indices in the UKBB. Testosterone (nmol/L), SHBG (nmol/L), and free androgen index (unitless) were observed to have varying distribution by sex, with outliers. Sex-specific extreme outliers for each phenotype were identified and excluded. Additionally, sex-specific normalization was performed to standardize each phenotype to mean 0 and SD 1 for analysis. SHBG= Sex-Hormone Binding Globulin; SD= Standard Deviation.

**Supplementary Figure 12.**
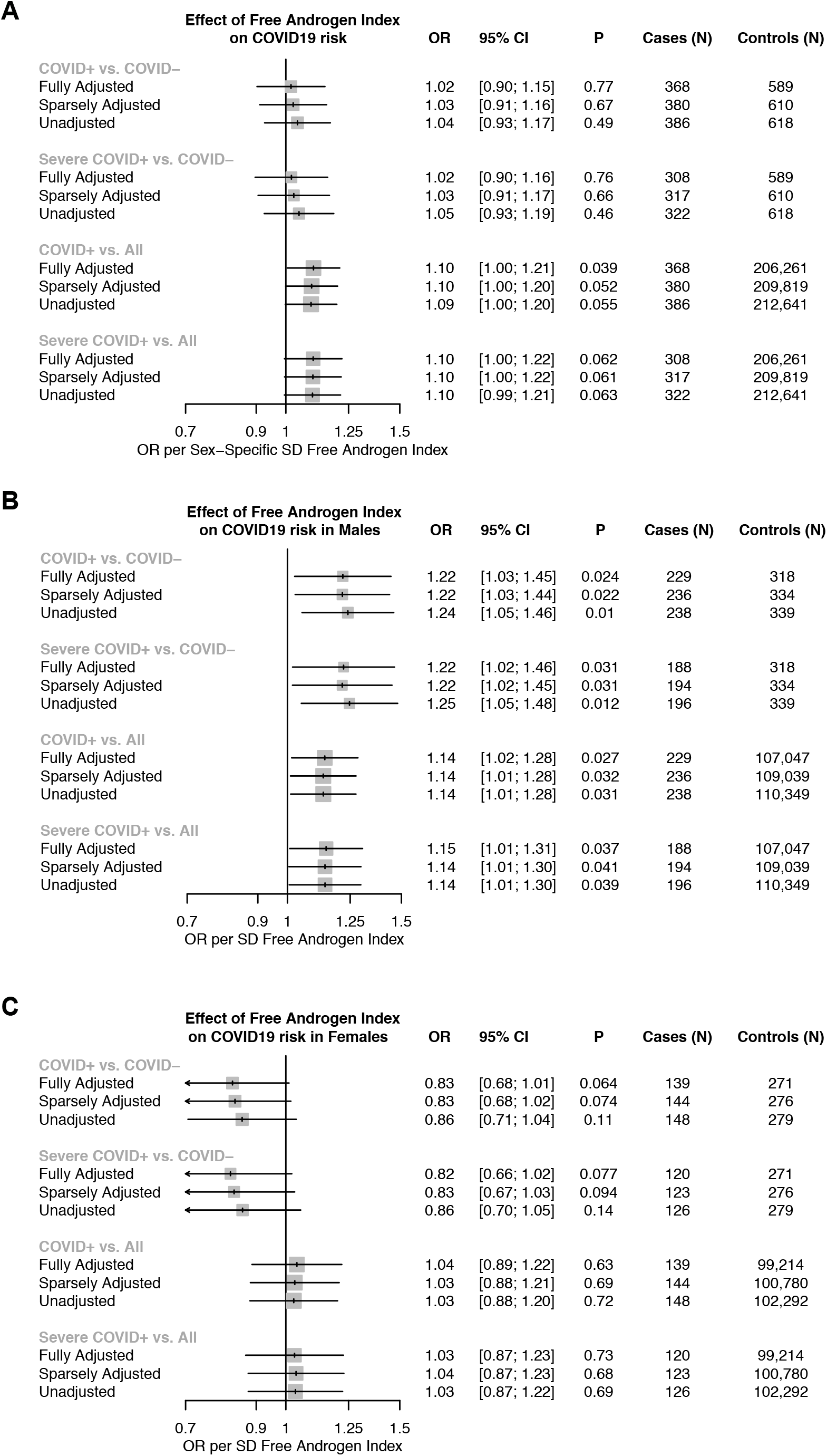
Association of free androgen index with Covid-19 in the UKBB dataset. Association of normalized free androgen index with four Covid-19 case/control models; for individuals tested with Covid-19: comparing 1) COVID+ vs. COVID-test results, 2) Severe COVID+ vs COVID-; and for all white British individuals from the UKBB English recruitment centers with available Covid-19 test reporting: comparing 3) COVID+ vs. all, 4) Severe COVID+ vs. all. Analyses are presented across three separate models: 1) unadjusted, 2) Sparsely adjusted for age, normalized Towndend deprivation index, and the first ten principal components of genetic ancestry, and 3) fully adjusted which additionally includes normalized body mass index as a covariate on top of the sparsely adjusted model covariates. Associations are displayed as **A.** Among males and females, **B.** among males, C. among females. COVID+= Tested positive for Coronavirus disease 2019, COVID-=Tested negative for Coronavirus disease 2019, all= all individuals with reported Covid-19 testing from the UKBB English recruitment centers., OR= Odds Ratio, SD= Standard Deviation, CI= Confidence Interval.

**Supplementary Figure 13.**
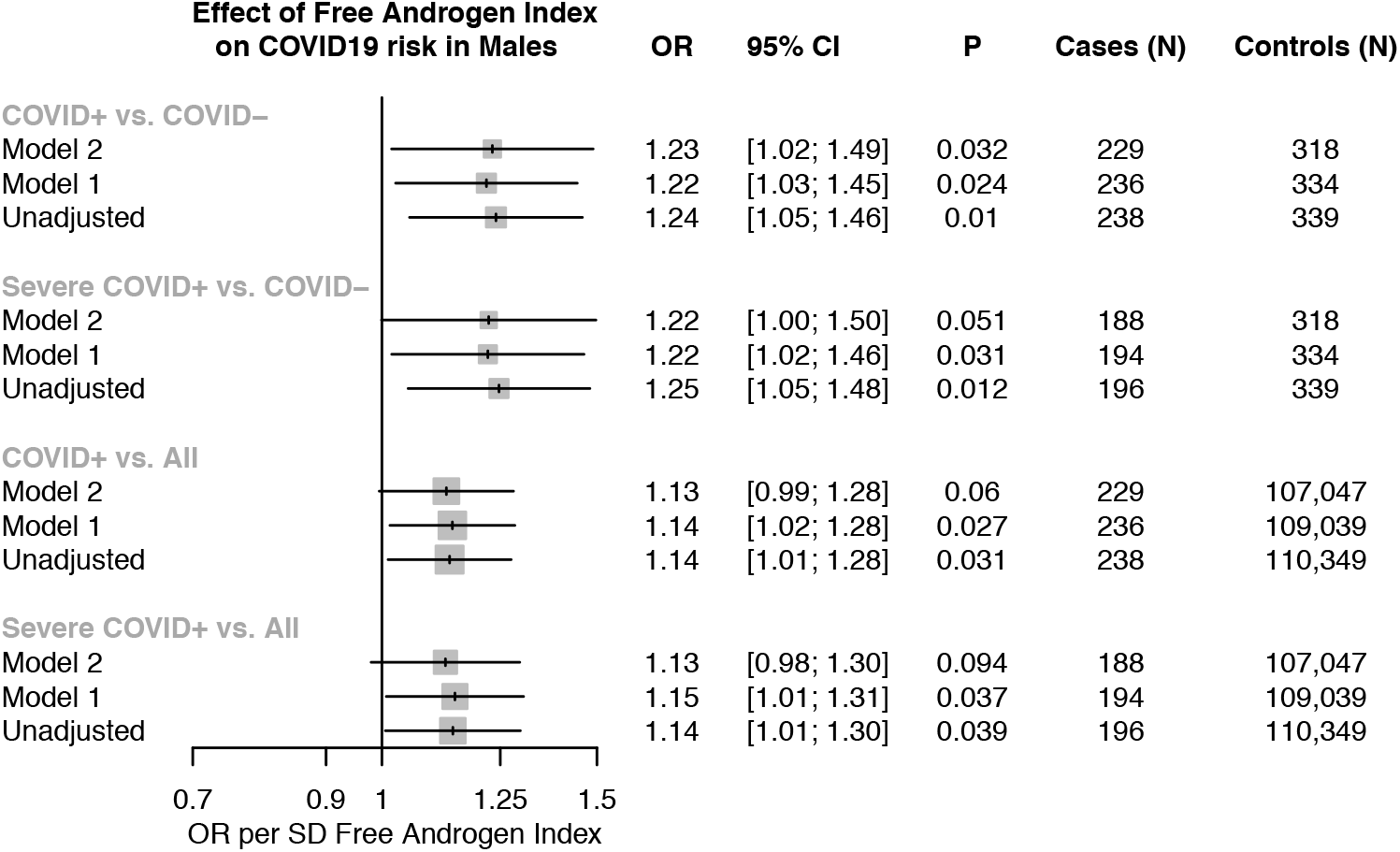
Adjusting for additional phenotypic covariates does not significantly influence the effect of free androgen index on COVID-19 risk. Association of normalized free androgen index in males across three separate adjustment models: 1) unadjusted, 2) Model 1 which is adjusted for age, normalized Townsend deprivation index, normalized body mass index, and the first ten principal components of genetic ancestry, and 3) Model 2 which additionally includes prevalent hypertension and diabetes as covariates on top of the sparsely adjusted model covariates. Analyses are presented across four COVID-19 case/control models: for individuals tested with COVID: comparing 1) COVID+ vs. COVID-test results, 2) Severe COVID+ vs. COVID-; and for all white British individuals from the UK Biobank English recruitment centers with available COVID-test reporting: comparing 3) COVID+ vs. All, 4) Severe COVID+ vs. All. COVID+ = Tested positive for coronavirus disease 2019, COVID-= Tested negative for coronavirus disease 2019, All = all individuals with reported COVID-19 testing from the UK Biobank English recruitment centers, OR = odds ratio, SD = standard deviation, CI= confidence interval.

**Supplementary Figure 14.**
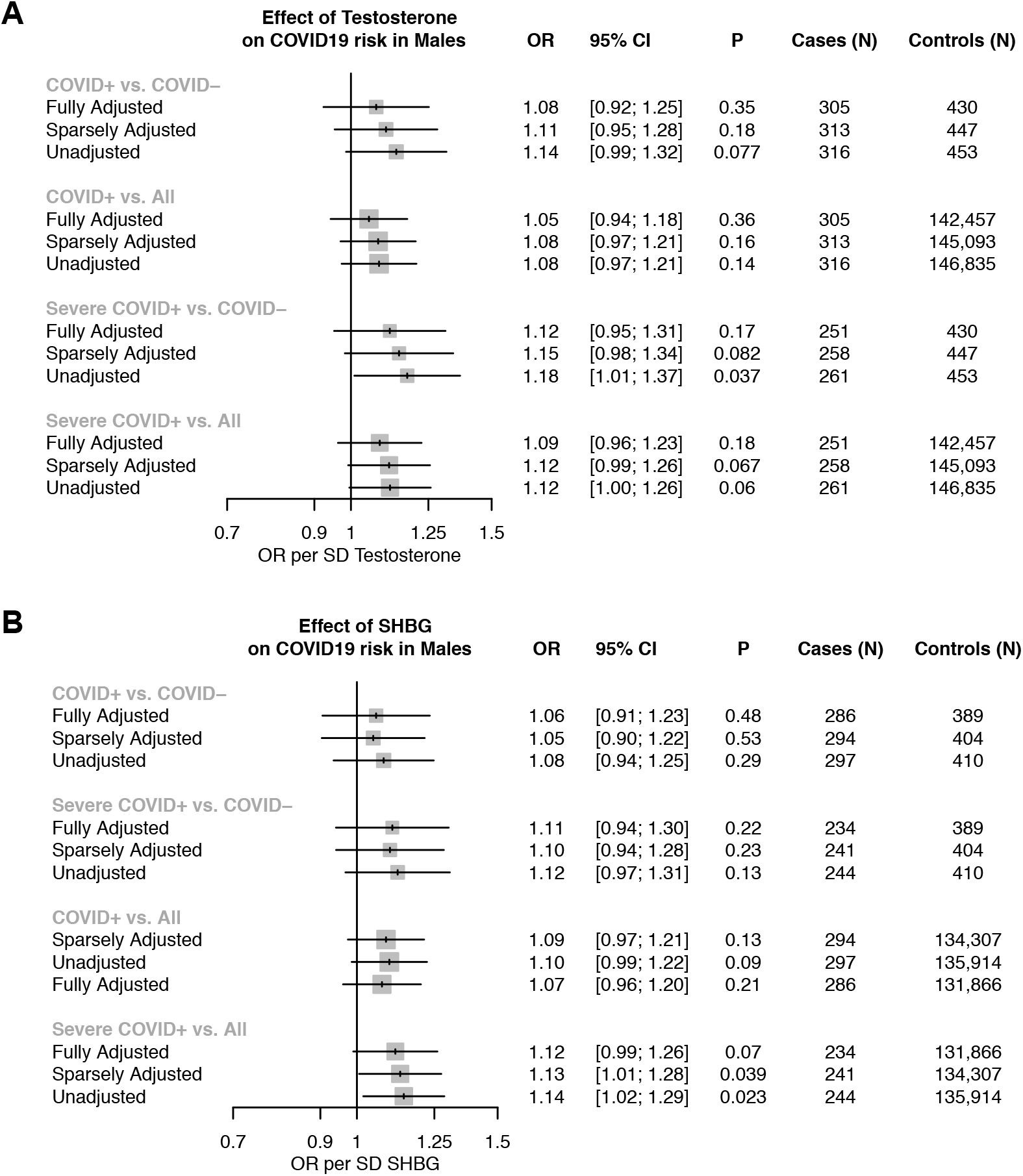
Association of testosterone and SHBG with COVID-19 in UK Biobank Males. Association of normalized testosterone and SHBG with four COVID-19 case/control outcome definitions: for individuals tested with COVID: comparing 1) COVID+ vs. COVID-test results, 2) Severe COVID+ vs. COVID-; and for all white British individuals from the UK Biobank English recruitment centers with available COVID-test reporting: comparing 3) COVID+ vs. All, 4) Severe COVID+ vs. All. Analyses are presented across three separate models: 1) unadjusted, 2) sparsely adjusted for age, normalized Townsend deprivation index, and the first ten principal components of genetic ancestry, and 3) fully adjusted which additionally includes normalized body mass index as a covariate on top of the sparsely adjusted model covariates. SHBG = sex-hormone binding globulin, COVID+ = Tested positive for coronavirus disease 2019, COVID-= Tested negative for coronavirus disease 2019, All = all individuals with reported COVID-19 testing from the UK Biobank English recruitment centers, OR = odds ratio, SD = standard deviation, CI= confidence interval.

## Supplementary Tables

**Supplementary Table 1.** Complete list of normalized z-scores for FDA drug library.

**Supplementary Table 2.** List of significantly predicted drug targets.

**Supplementary Table 3.** List of genes in “GO AR signaling”, “Upstream AR activators” and “AR target genes” expression modules.

**Supplementary Table 4.** List of phenotypic covariates included in sensitivity analysis.

